# A Bacterial Natural Product Reshapes Phyllosphere Microbiome Composition by Blocking Carotenoid Biosynthesis

**DOI:** 10.64898/2026.05.14.725204

**Authors:** Mike Wallace, Madelyn L. Allen, Talia L. Karasov, Aaron W. Puri

## Abstract

The microbial communities that inhabit the phyllosphere, the above-ground portion of plants, are important for plant growth and resilience. However, the natural products that mediate interactions among these organisms and with their environment remain understudied, limiting our molecular-level understanding of how phyllosphere microbial communities are structured and maintained. Natural product-mediated microbial competition is typically associated with growth inhibition, but may also occur through other mechanisms in these communities. Many bacteria produce pigments to mitigate oxidative stress, and here we identify listianol, a previously undescribed natural product that inhibits the pigmentation of diverse bacteria. Listianol blocks carotenoid biosynthesis by targeting the desaturases CrtN and CrtI. This activity sensitizes normally pigmented bacteria to UVB radiation *in vitro*, and a listianol-producing strain reshapes the composition of a model bacterial community on *Arabidopsis thaliana* under UVB exposure. Together, these findings identify pigmentation inhibition as a potential form of competition in the phyllosphere.

## INTRODUCTION

The phyllosphere, or aerial plant surface, hosts bacterial communities with complex relationships both among themselves and with their plant hosts. These communities can strongly influence plant health, including by stimulating plant growth and providing protection from biotic and abiotic stresses^1,2^. Bacterial constituents of the phyllosphere are exposed to reactive oxygen species (ROS) generated by the plant host in response to stresses, including drought, infection, or phytotoxic compounds, as well as from abiotic sources such as heavy metals and UV light^3,4^. To protect themselves from ROS, many phyllosphere bacteria produce ROS-quenching pigments, including arylpolyenes and carotenoids^1,3^.

Pink-pigmented facultative methylotrophs (PPFMs) are common constituents of the phyllosphere microbiota^1,5^. These bacteria are metabolically versatile and can use reduced one-carbon compounds as their sole source of carbon and energy^6^. This includes methanol, which is a byproduct of plant carbon cycling and is released from stomata^7,8^. In some instances, PPFMs have been shown to improve the growth of plants, possibly through the production of stimulatory plant hormones such as auxin and cytokinins^9–11^. Recently, there is growing industrial interest in using PPFMs as bioinoculants to protect crops from pests and abiotic stress^12,13^. However, little is known about how PPFMs interact with other phyllosphere microorganisms. This limits researchers’ abilities to take advantage of beneficial interactions and to understand and predict the phyllosphere microbiota structure under different environmental conditions.

Many bacterial interactions are mediated by natural products which can facilitate cooperation and competition with other microbes as well as the plant host. For example, natural products are often used to gain a competitive advantage by enabling the sequestration and uptake of nutrients such as iron^14^, or by inhibiting the growth of other strains that occupy the same niche^15^. In bacteria, the biosynthetic enzymes required to produce natural products are often encoded by co-localized genes within the genome as biosynthetic gene clusters (BGCs). BGC prediction tools suggest that PPFMs may have significant and underexplored biosynthetic potential^16,17^. However, few unique natural products have been isolated from PPFMs to date^18,19^.

In many cases BGCs are transcriptionally silent under lab conditions^20^, which impedes our ability to characterize the structure and function of the natural products made by these gene clusters. One method for activating silent BGCs is heterologous expression. A promising platform for the expression of silent BGCs is chassis-independent recombinase-assisted genomic engineering (CRAGE)^21^. The CRAGE system simplifies the traditional heterologous expression workflow by streamlining the introduction of heterologous BGCs and by improving the stability of BGC expression via genome integration.

In this work, we develop a heterologous expression platform for exploring the biosynthetic potential of PPFMs based on the CRAGE system. Using this platform, we discover a new PPFM natural product, listianol. We find that listianol blocks carotenoid production in diverse bacteria by inhibiting the desaturase enzymes involved in their biosynthesis, which is, to our knowledge, a new mechanism of action for a natural product. We show that through inhibiting bacterial pigmentation, bacteria treated with listianol are more sensitive to UV radiation *in vitro*, and that a listianol-producing strain alters the composition of a synthetic bacterial community in *Arabidopsis thaliana* under UV exposure. These results identify a new PPFM natural product that provides a link between inhibition of pigment biosynthesis and ecological competition in the phyllosphere.

## RESULTS

### Development of a heterologous expression platform for BGCs from PPFMs

In order to explore the biosynthetic potential of PPFMs, we chose a modified variant of the well-studied PPFM strain *Methylorubrum extorquens* PA1 as our heterologous expression host^22^. This strain has deletions that reduce flocculation in liquid culture^23^ and that eliminate the production of two acyl-HSL quorum sensing signals^24^, facilitating easier growth and isolation of any newly produced natural products. We first introduced the genome landing pad (LP) for BGCs via transposon mutagenesis, and selected a mutant, PA1-LP, where the LP had been inserted into a YvrE-type sugar lactonase. PA1-LP had no growth defects or other obvious phenotypic changes (Figure S1).

The original CRAGE platform uses the inducible Lac-T7 expression system^25^ to regulate expression of inserted BGCs with isopropyl β-D-1-thiogalactopyranoside (IPTG)^21^. Unfortunately, we found that this system worked poorly in *M. extorquens* PA1 (Figure S2). To address this, we assembled several IPTG-inducible *lux* reporter plasmids with different promoters, including P_L/O4/A1_, a promoter developed to optimize gene expression in *Methylorubrum extorquens* AM1, a closely related strain to PA1^26^. When integrated into PA1-LP, all new promoters exhibited increased luminescence over P_T7_, with P_L/O4/A1_ exhibiting ∼30-fold higher luminescence and greater fold change upon induction (Figure S2). We therefore chose the P_L/O4/A1_ promoter to drive expression of BGCs in PA1-LP.

We next selected BGCs for heterologous expression by examining clusters identified using the tool antiSMASH^27^ in publicly available genomes of *Methylorubrum* and *Methylobacterium*. For this initial study, we focused on BGCs that were likely to produce novel natural products, rather than highly conserved clusters. We selected BGCs that were co-directional, of a size that is practical to assemble, and that have plausible boundaries that can be inferred via analysis of the genome neighborhood surrounding the core biosynthetic genes. These criteria led us to select seven BGCs to screen via heterologous expression in this initial work (Table S1). These BGCs were assembled under P_L/O4/A1_ and integrated into PA1-LP to generate seven different strains expressing the chosen BGCs.

To identify new metabolites, we cultured, extracted and analyzed each strain by reverse-phase liquid chromatography high-resolution tandem mass spectrometry (LC-HRMS/MS), using both HRMS/MS and UV-Vis readouts for analysis. Of the seven BGCs screened, three produced metabolites with detectable UV absorbance: BGC2, BGC4, and BGC7 (Figure 1a). Given the intensity of the features produced by the strain expressing BGC4, we chose to further characterize its products.

**Figure 1.**
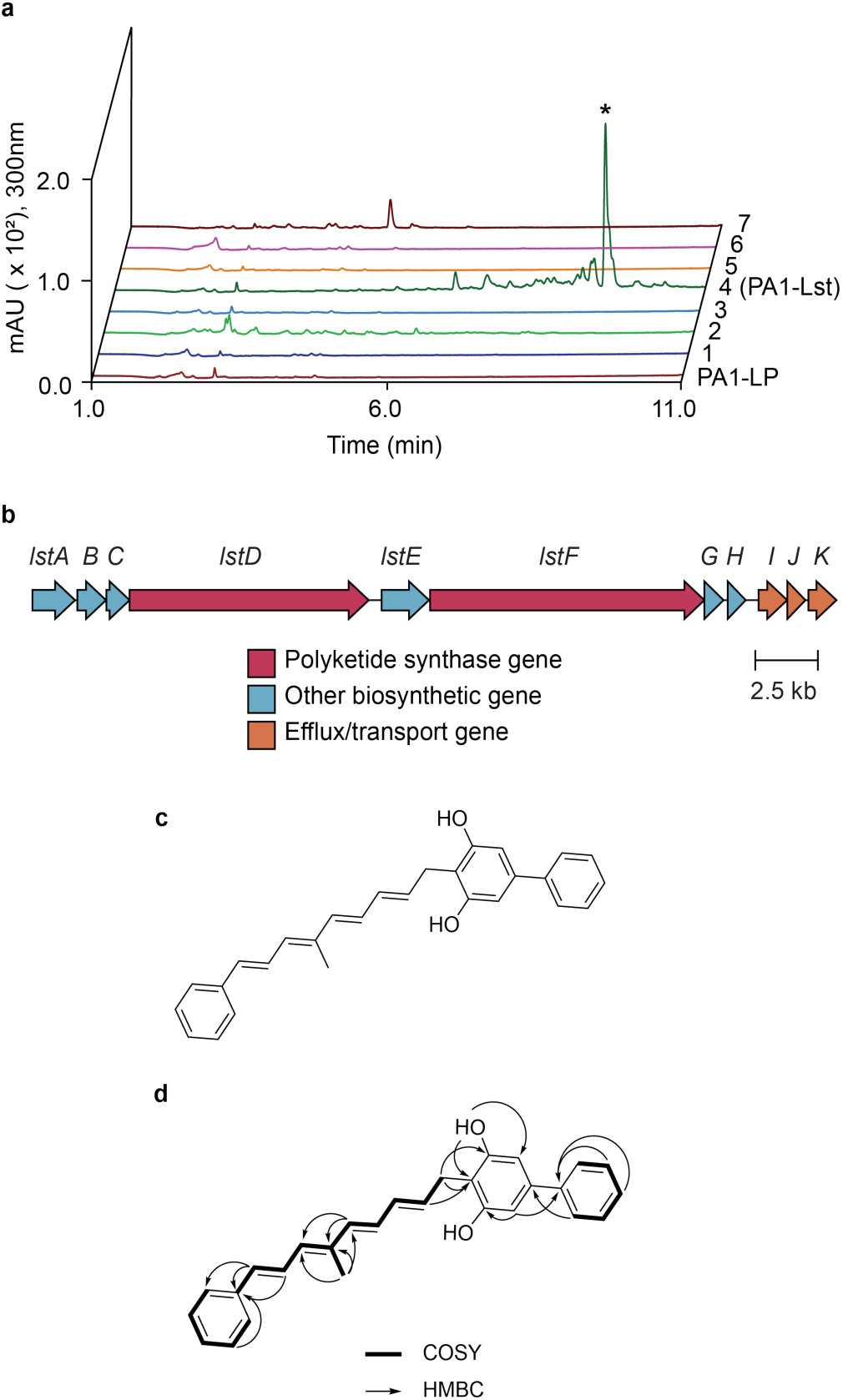
A screen of BGCs from PPFMs uncovers new metabolites, including the diarylresorcinol listianol. (**a**) Analysis of extracts of PA1-LP strains harboring BGCs 1-7, displayed as UV-Vis chromatograms extracted at 300nm. The asterisk indicates the UV-Vis feature corresponding to listianol. (**b**) Genetic organization of the *lst* BGC, BGC4. (**c**) Structure of listianol, the isolated metabolite from PA1-Lst. (**d**) Structure of listianol with key ^1^H-^1^H COSY and ^1^H-^13^C HMBC correlations shown.

### BGC4 produces listianol, a new diarylresorcinol natural product

BGC4 is from the genome of the PPFM *Methylobacterium* sp. 4-46 and is predicted to encode a hybrid resorcinol/polyketide synthase (PKS) BGC (Figure 1b). We only found this BGC in one other strain, the PPFM *Methylobacterium* sp. WSM2598. Notably, both strains were isolated in Africa from the flowering legume *Listia bainesii* by separate research groups decades apart^28–30^. We chose to focus on the most abundant metabolite produced by BGC4 for isolation and structural characterization. HRMS analysis of this product gave a presumed [M-H]^-^ *m/z* value of 393.1867, corresponding to a molecular formula of C_28_H_26_O_2_ and 16 degrees of unsaturation. Consistent with the high degree of unsaturation, this metabolite exhibited a maximum absorbance at 349nm with shoulders at 333nm and 366nm, indicating the presence of a polyene moiety (Figure S3). The UV-Vis absorbance spectrum contained an additional maximum at 260nm, possibly indicating the presence of aromatic group(s). We solved the structure of this metabolite using 1– and 2-dimensional NMR experiments as well as HRMS/MS (Figures S4-S8, Table S2). The structure of the primary product of BGC4 is a highly unsaturated diarylresorcinol natural product (Figures 1c and 1d). Because both strains known to carry this BGC were isolated from *Listia bainesii,* as well as the presence of the resorcinol moiety, we named this metabolite listianol. Lastly, to determine if we had omitted any important genes from this cluster, we assembled an extended version of BGC4 that included additional downstream genes encoding another predicted thioesterase as well as an ABC-type multidrug transport system. Upon integration into PA1-LP, we found this new strain, which we refer to as PA1-Lst, also produced listianol (Figure S9). PA1-Lst was used for all further experiments.

Canonically, disubstituted resorcinol-containing natural products are synthesized by a DarB-type ketosynthase and a DarA-type aromatase^31,32^. The *lst* BGC is predicted to encode a DarB-type ketosynthase (LstB) and a DarA-type aromatase (LstC), as well as two PKSs (LstD and LstF). The BGC also encodes a predicted aromatic amino acid ammonia-lyase (LstA), which could generate cinnamic acid from phenylalanine. Based on the structure of listianol and the predicted enzymes encoded by the *lst* BGC, we propose a plausible biosynthesis for this molecule that involves condensation of a cinnamic acid-derived β-ketoacyl polyketide product and a second molecule of cinnamic acid (Figure 2). To test this hypothesis, we constructed Δ*lstB* and Δ*lstC* mutants and detected the expected intermediates (Figure S10). We also detected incorporation of the predicted number of carbons from phenylalanine and cinnamic acid into listianol and the intermediates using inverse stable isotopic labeling (Figures S10 and S11). Together, these results support our proposed biosynthesis of listianol.

**Figure 2.**
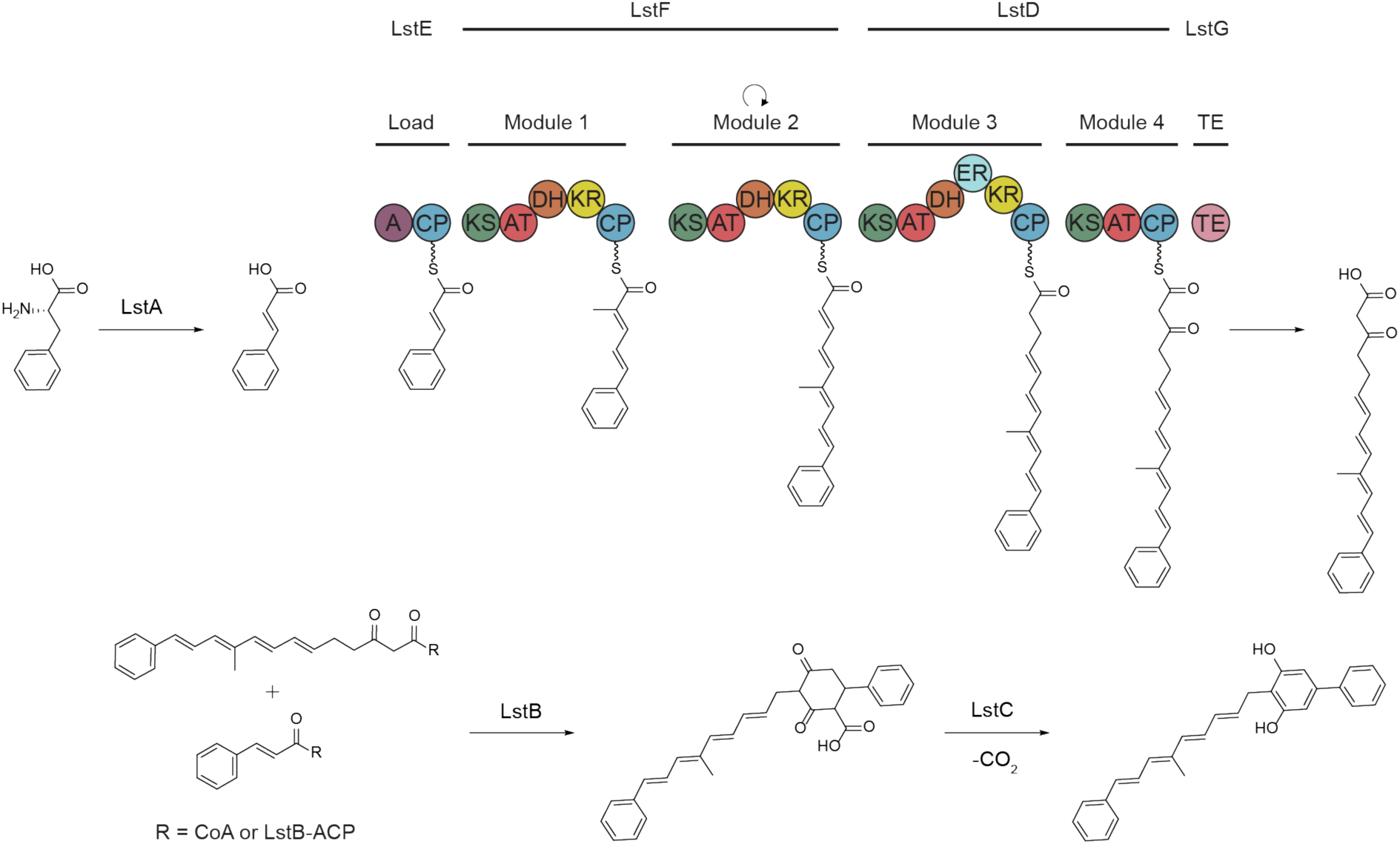
Proposed biosynthesis of listianol. Polyketide synthase (PKS) domains are shown as circles within each predicted PKS module. Letters indicate adenylation (A), acyl carrier protein (CP), ketosynthase (KS), acyltransferase (AT), dehydratase (DH), ketoreductase (KR) and thioesterase (TE) domains. The curved arrow indicates a predicted repeated condensation during biosynthesis.

### Listianol prevents carotenoid biosynthesis via inhibition of the desaturase enzymes in diverse bacteria

We noticed a significant difference in the pigmentation of PA1-Lst compared to the base CRAGE strain, PA1-LP. While PA1-LP colonies are pink, PA1-Lst colonies are white. To further investigate, we grew PA1-LP, as well as the taxonomically distant gram-positive bacterium *Cellulomonas* sp. Leaf334, with the addition of purified listianol, which also resulted in unpigmented cultures (Figure 3a). LC-HRMS/MS analysis of the extracted cell pellets showed the disappearance of products that, based on UV-Vis absorbance spectra, are likely carotenoids (Figure S12). We quantified the inhibition of carotenoid biosynthesis using whole cell assays and found that listianol was considerably more potent against PPFMs compared to other strains tested, with IC_50_ values in the low nanomolar range (Figure 3b, Table S3).

**Figure 3.**
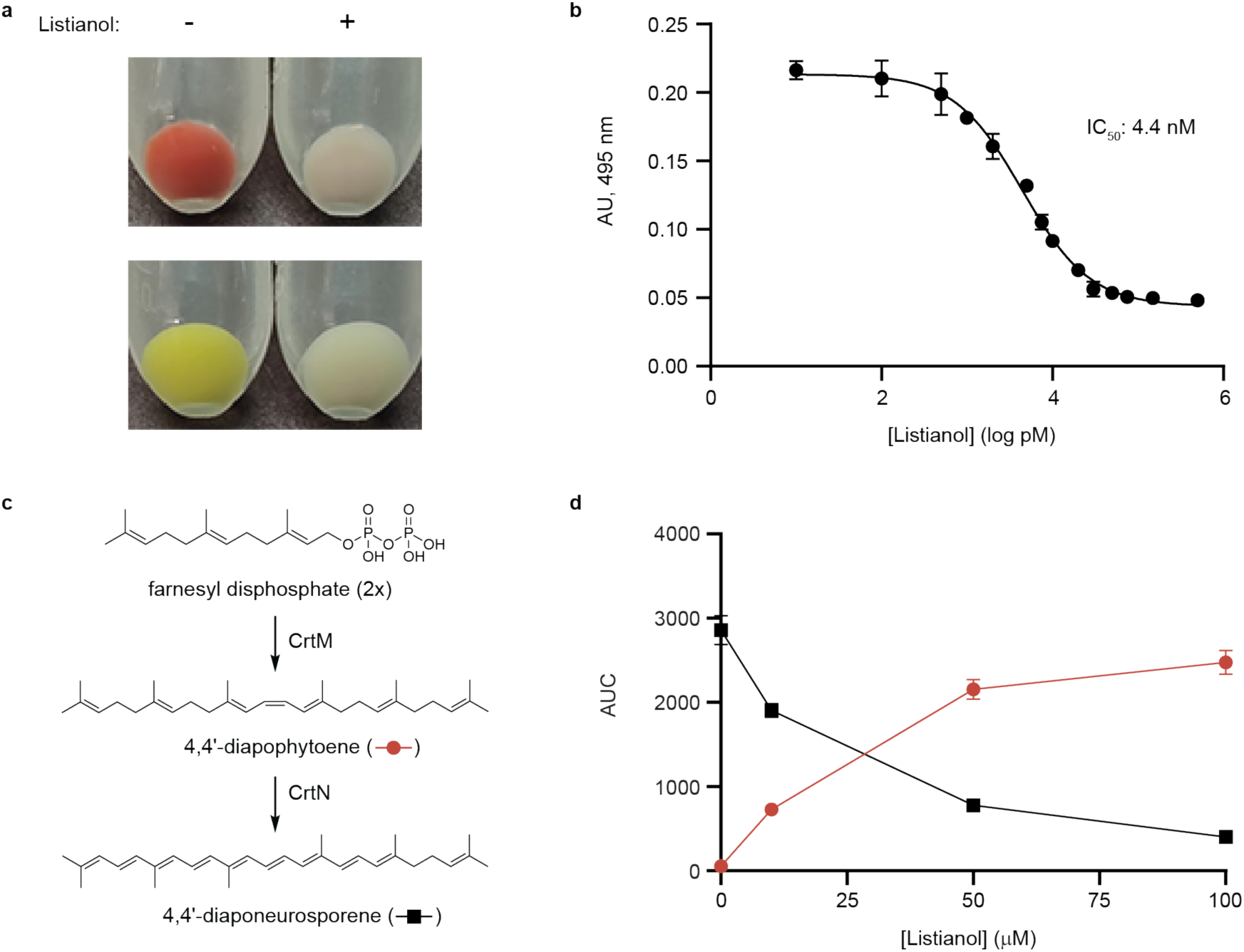
Listianol inhibits carotenoid biosynthesis in diverse bacteria by inhibiting the desaturase enzyme CrtN. (**a**) Cell pellets from cultures of PA1-LP (top) and *Cellulomonas* sp. Leaf334 (bottom) grown in the absence or presence of 1 µM listianol. (**b**) Quantified inhibition of carotenoid biosynthesis in PA1-LP cultures grown with exogenous listianol. AU, absorbance units. (**c**) Scheme illustrating the first steps of C30-carotenoid biosynthesis. (**d**) Dose-dependent inhibition of CrtN in *E. coli* – *crtMN* by listianol results in disappearance of 4,4’-diaponeurosporene (red circles) and accumulation of 4,4’-diapophytoene (black squares). The y-axis represents the area under the curve (AUC) for each product when analyzed by HPLC. CrtM products were extracted at 288nm and the CrtMN products at 450nm. Error bars indicate standard deviation of n=2 biological replicates.

Both PPFMs and *Cellulomonas* sp. Leaf334 are pigmented due to carotenoids, metabolites that are produced by many diverse organisms^33,34^. This led us to hypothesize that listianol may prevent pigment formation through inhibition of carotenoid biosynthesis. Carotenoids are isoprene-derived metabolites that contain a highly conjugated polyene chain, often with the presence of additional functional groups that vary depending on the organism^35^. They are categorized according to carbon chain length, with C30 and C40 carotenoids representing the most prevalent classes^33^. A well characterized C30-carotenoid is staphyloxanthin, produced by methicillin-resistant *Staphylococcus aureus* (MRSA) clinical isolates^36,37^. Biosynthesis of staphyloxanthin begins with the condensation of two farnesyl diphosphate molecules to form 4,4’-diapophytoene by the enzyme 4,4’-diapophytoene synthase, CrtM. 4,4’-diapophytoene is then oxidized to form the conjugated polyene 4,4’-diaponeurosporene by the enzyme 4,4’-diapophytoene desaturase, CrtN^38^ (Figure 3c). Subsequently, 4,4’-diaponeurosporene is further modified to form the final carotenoid staphyloxanthin. Biosynthesis of C40-carotenoids begins with the same chemical transformations but instead uses geranylgeranyl diphosphate to form phytoene and lycopene, via the enzymes CrtB and CrtI, respectively^33^. PPFMs also produce C30-carotenoids, but the enzymes HpnC-E form squalene from farnesyl diphosphate, which is then oxidized by CrtN^33,39^.

To determine if listianol inhibits carotenoid biosynthesis, we heterologously expressed CrtM and CrtM + CrtN from *S. aureus* in *E. coli.* We found that *E. coli* – *crtM* produced a metabolite with an absorbance spectrum consistent with that of 4,4’-diapophytoene and that *E. coli* – *crtMN* produced a metabolite with an absorbance spectrum consistent with that of 4,4’-diaponeurosporene (Figures S13 and S14). When we grew *E. coli – crtM* with listianol, the abundance of 4,4’-diapophytoene was unchanged (Figure S15a). Conversely, when we grew *E. coli* – *crtMN* with listianol the abundance of 4,4’-diaponeurosporene decreased, while the abundance of 4,4’-diapophytoene increased (Figure 3d), consistent with listianol inhibiting the 4,4’-diapophytoene desaturase CrtN. As a positive control, we also tested these same strains with a known inhibitor of CrtN, the synthetic antifungal naftifine^37^, and observed the same decrease in abundance of 4,4’-diaponeurosporene and accumulation of the 4,4’-diapophytoene intermediate (Figures S15a and S15b). We found similar results when we heterologously expressed the analogous enzymes in the C40-carotenoid biosynthetic pathway, CrtB and CrtBI, from *Erwinia herbicola*^40^ (Figures S13-S15c-f).

While performing these assays, we also noticed that listianol inhibited the growth of some bacterial strains at higher concentrations (Table S3). Of note, listianol was effective against the human pathogen MRSA at 10 μg mL^-1^ and the plant pathogens *Clavibacter michiganensis* ssp. *michiganensis* and *Curtobacterium flaccumfaciens* at 5 and 10 μg mL^-1^, respectively. We do not know the mechanism of action for this growth inhibition, however it is likely unrelated to carotenoid inhibition, as the growth of both pigmented and unpigmented strains was inhibited by listianol. Together these results indicate that listianol is a multifunctional natural product that inhibits the growth of bacteria and carotenoid biosynthesis, likely via inhibition of the desaturase enzymes CrtN and CrtI.

### Listianol sensitizes pigmented bacteria to UVB radiation *in vitro*

Given that PPFMs are primarily epiphytic and would be regularly exposed to UV light, we hypothesized that listianol may alter the phyllosphere microbiota composition by sensitizing these organisms to UV-induced ROS via inhibition of carotenoid biosynthesis. To test this hypothesis, we determined the UVB survivability of cultures of *Cellulomonas* sp. Leaf334, *Methylobacterium* sp. Leaf456 and PA1-LP grown in the absence or presence of listianol and found that the listianol-treated cultures were significantly more susceptible to UVB radiation (Figure 4a). We also found that the PPFMs were markedly more resistant to UVB than *Cellulomonas* sp. Leaf334, even when pigmentation was inhibited, suggesting that these bacteria have other UVB protection mechanisms in addition to carotenoids to protect themselves. Together, these results demonstrate that listianol sensitizes pigmented bacteria to UVB radiation.

**Figure 4.**
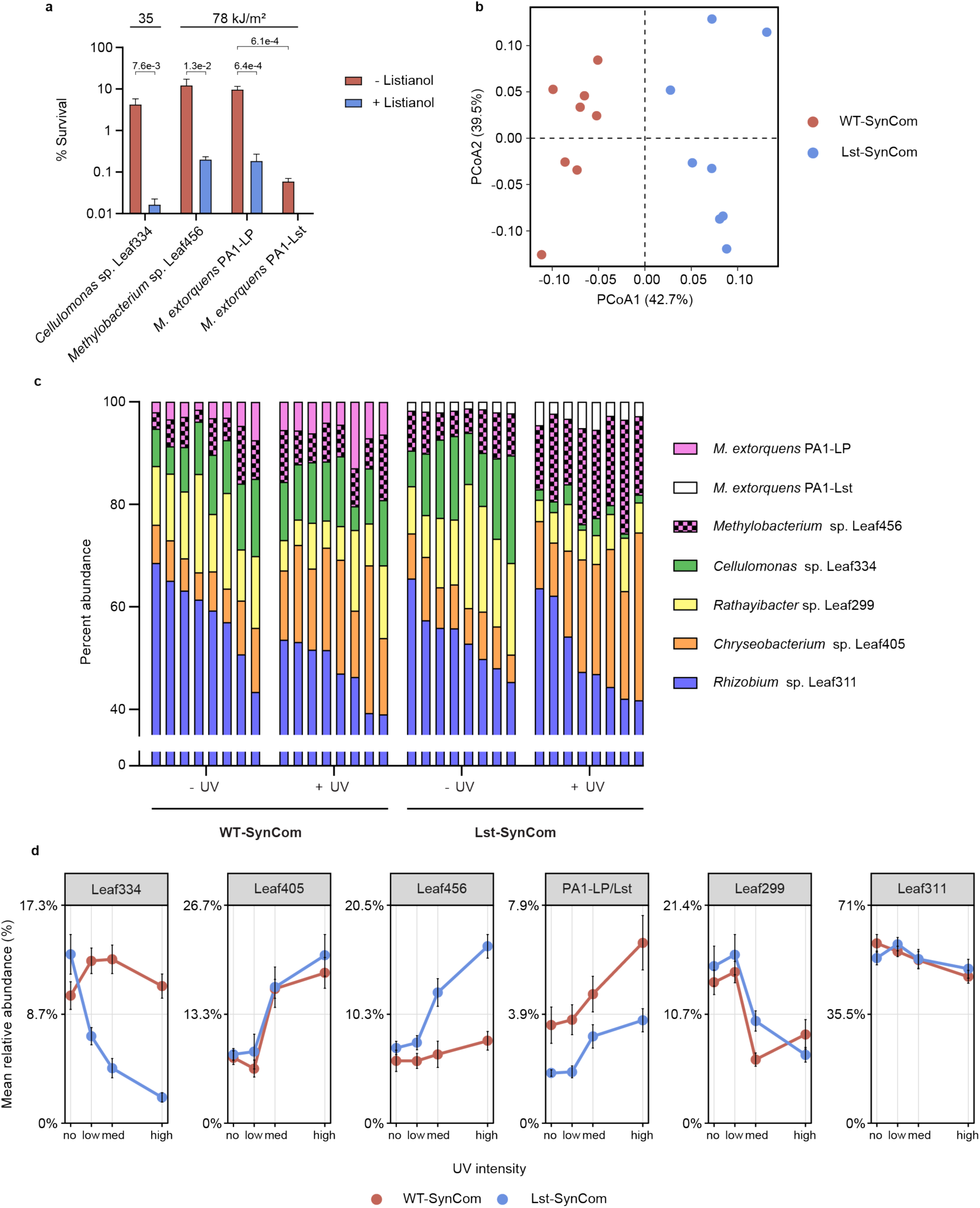
Listianol alters the composition of an *Arabidopsis thaliana* phyllosphere SynCom when exposed to UV. (**a**) UV susceptibility of indicated SynCom strains when grown with listianol *in vitro*. The UV dose used to test each strain are shown by the numbers at the top of the bar plots. The error bars indicate standard deviation and statistical differences were determined by two-tailed unpaired t-test (n=3). Brackets above the bar plots indicate comparison groups and the numbers indicate *p*-values. (**b**) PCoA plot of Bray-Curtis dissimilarities between bacteria recovered from plants inoculated with WT-SynCom and Lst-SynCom and grown under high-UV irradiation. Each circle represents the bacterial community isolated from a single plant. (**c**) Relative abundances of SynCom constituents. Each stacked bar represents the bacterial community isolated from a single plant. The UV data shown here is from one replicate of the high-UV condition. (**d**) The mean relative abundance for each strain in either the WT – or Lst-SynCom, across all UV conditions. The x-axis represents the intensity of UVB radiation to which the plants were exposed. Low corresponds to 0.5 W m^-2^, med to 1.0 W m^-2^ and high to 2.2 W m^-2^. Error bars represent standard error (n=8).

### Listianol alters the composition of a synthetic phyllosphere community of bacteria under UVB stress on *Arabidopsis thaliana*

Finally, to determine if listianol has activity *in planta*, we assembled a small synthetic community (SynCom) of phyllosphere strains from the *At*-LSPHERE strain collection^41^, a group of bacterial strains isolated from the phyllosphere of wild *Arabidopsis thaliana* plants, and explored the impact of listianol on this community under UV stress on *A. thaliana*. To assemble the SynCom, we selected five strains shown to be strong colonizers of *A. thaliana* ^42,43^ that represented different possible combinations of sensitivity to pigmentation and growth inhibition by listianol. In addition to carotenoid-pigmented strains, we also included an unpigmented strain and an aryl polyene-producing strain, to represent the other major pigment class found in phyllosphere bacteria. Finally, we added either the strain PA1-LP as a control or PA1-Lst to form WT-SynCom and Lst-SynCom, respectively, for inoculation of *A. thaliana* (Table 1).

**Table 1.** SynCom members and their susceptibility to listianol.

| Strain | Pigmentation Type | Carotenoid Inhibition | Growth Inhibition |
| --- | --- | --- | --- |
| *Cellulomonas* sp. Leaf334 | Carotenoid | Yes | 5 µg mL<sup>-1</sup> |
| *Chryseobacterium* sp. Leaf405 | Aryl polyene | Not applicable | 40 µg mL<sup>-1</sup> |
| *Methylobacterium* sp. Leaf456 | Carotenoid | Yes | No |
| *Rathayibacter* sp. Leaf299 | Carotenoid | No | 10 µg mL<sup>-1</sup> |
| *Rhizobium* sp. Leaf311 | Unpigmented | Not applicable | No |

We inoculated *A. thaliana* seedlings with WT-SynCom or Lst-SynCom and grew these plants for four days under no UV stress, followed by six days of exposure to UVB radiation at three different environmentally relevant intensities. The three intensities, low-UV, med-UV, and high-UV, represent average daily UV doses received in Germany (low-UV), where many of the *At-*LSPHERE strains originate, and low-altitude South Africa, where *Methylobacterium* sp. 4-46 originates, during moderate (med-UV) or intense UV conditions (high-UV)^41,44–46^. We then harvested the plants and determined the abundance of each strain in the SynComs under the different UV conditions.

We found that the presence of the listianol-producing strain had a distinct effect on the composition of the SynCom compared to the control (Figure 4b). Specifically, in the presence of UV, *Cellulomonas* sp. Leaf334 exhibited a statistically significant decrease in relative abundance, but only when PA1-Lst was present (Figure 4c). The change was significant even in the low-UV condition but became much more apparent under the higher UV intensities (Figure 4d). This is consistent with listianol sensitizing *Cellulomonas* sp. Leaf334 to UV radiation *in planta*, as it did *in vitro* (Figure 4a).

In addition to changes in the relative abundance of *Cellulomonas* sp. Leaf334, the abundance of other community members changed under UV stress, but independent of the presence listianol (Figure 4d). *Rhizobium* sp. Leaf311 and *Rathayibacter* sp. Leaf299 both decreased in abundance as UV intensity increased, while *Chryseobacterium* sp. Leaf405 and the PPFMs increased in abundance. As *Cellulomonas* sp. Leaf334 decreases in relative abundance in the presence of UV and listianol, different strains filled this vacancy in different replicates of the experiment. This included the PPFM *Methylobacterium* sp. Leaf456 as well as *Chryseobacterium* sp. Leaf405 and *Rhizobium* sp. Leaf311 (Figure 4d, Figure S16). This shows no other SynCom strain consistently benefited from the presence of listianol in this system.

While listianol shows a significant effect on the bacterial composition of the SynCom, it does not appear to impact the plant itself. We observed decreases in plant weight for plants grown under increasing UVB, but the presence of PA1-Lst in the SynCom had no effect on plant weight or pigmentation (Figures S17 and S18). Additionally, there were no differences in plant health or pigmentation when inoculated with PA1-Lst alone, indicating that listianol does not have a significant impact on plant health (Figure S19). Together, these results illustrate that listianol can alter the composition of a phyllosphere SynCom in *A. thaliana* under UV radiation.

## DISCUSSION

PPFMs are common members of the phyllosphere microbiota with significant biosynthetic potential, but little is known about the natural products these organisms produce or their functions in the phyllosphere. Here, we developed a platform to characterize PPFM natural products based on the CRAGE system. Using this platform, we discovered listianol, a unique inhibitor of the desaturase enzymes involved in carotenoid biosynthesis, CrtN and CrtI. We then demonstrate that listianol sensitizes certain carotenoid-producing bacteria to UV radiation *in vitro* and alters an *A. thaliana* phyllosphere SynCom in the presence of UV radiation, thereby identifying pigmentation inhibition as a potential form of competition in the phyllosphere.

To our knowledge, listianol is the first bacterial natural product known to directly inhibit carotenoid biosynthesis via inhibition of the desaturases CrtN and CrtI. There are examples of fungal and bacterial natural products, including mevinolin^47^ and fosmidomycin^48^, that block carotenoid biosynthesis, but they do so indirectly by inhibiting the biosynthesis of carotenoid precursors. Given that the production of 4,4’-diapophytoene and phytoene are not inhibited by treatment with listianol (Figures 3d and S15), it is unlikely that listianol inhibits the enzymes required for production of the isoprene precursors in carotenoid biosynthesis. While the listianol BGC was only found in the genomes of two PPFMs, given the abundance of ROS-quenching pigments amongst phyllosphere bacteria and their importance in survival^3,49,50^, it is possible that natural products that inhibit pigment biosynthesis are more common than previously appreciated.

We have shown that a listianol-producing strain alters the composition of a phyllosphere SynCom in the *A. thaliana* model system when exposed to UV radiation. This is consistent with listianol sensitizing pigmented strains to UV-induced oxidative damage via carotenoid biosynthesis inhibition. Specifically, we observed a UV – and listianol-dependent decrease in the abundance of *Cellulomonas* sp. Leaf334. We cannot ignore the antimicrobial effects of listianol against *Cellulomonas* sp. Leaf334 as a cause for this decrease (Table 1), however we think this is unlikely because the abundance of this strain only decreases in the presence of UV exposure. If the antimicrobial activity were responsible, we would expect this effect to also be present in the absence of UV. There are several examples of UV light altering the microbiota composition of plants and algae, as well as carotenoid-deficient mutants exhibiting reduced populations in field experiments^49–52^.

Extrapolating these findings to an environmental setting is challenging given the differences between laboratory conditions and native environments. Additionally, because listianol was discovered via heterologous expression, we do not know under what conditions the listianol BGC is expressed in the native producer *Methylobacterium* sp. 4-46, nor if/how this strain protects itself from the effects of listianol. Presumably, listianol enables the native strain to gain a competitive advantage in its natural environment, possibly through sensitizing other bacteria to UV radiation or other sources of ROS. Listianol most potently inhibits carotenoid biosynthesis in PPFMs (Table S3), yet we found that even unpigmented PPFMs are less susceptible to lower levels of UV radiation than another phyllosphere strain, *Cellulomonas* sp. Leaf334 (Figure 4a). At these UV levels, PPFMs may not need carotenoids to survive, instead relying on alternative mechanisms to cope with UV-induced ROS stress^19,53^, while other phyllosphere strains would be sensitized to UV by listianol. At higher UV levels, because listianol still results in increased sensitivity to UV radiation in PPFMs, it may aid the native producer in competing against other PPFMs. At these UV levels, the native producer would likely need a mechanism to protect itself from the inhibitory effects of listianol, such as enhanced listianol efflux or upregulation of ROS-quenching enzymes and DNA repair pathways.

The PPFM CRAGE system developed here illustrates the biosynthetic potential PPFMs hold for natural products research. Interestingly, all the expressed BGCs that produced new metabolites were PKS-based, while the predicted beta-lactone and nonribosomal peptide synthetase BGCs failed to produce any detectable metabolites. This could represent a limitation of *M. extorquens* PA1 as a host for heterologous expression of BGCs from these natural product classes or a need for better predictions of BGC boundaries. Further research will be needed to uncover the new metabolites produced by the other BGCs identified in this work.

In summary, we have discovered a natural product originating from PPFMs that inhibits bacterial carotenoid biosynthesis. Listianol can sensitize carotenoid-producing bacteria to UV radiation and alters the composition of an *A. thaliana* phyllosphere SynCom in the presence of UV, thereby establishing a link between inhibition of pigment biosynthesis and ecological competition in the phyllosphere. This study also demonstrates the potential that PPFMs hold for the discovery of natural products with unique structures and functions. Through the discovery of natural products made by members of the phyllosphere microbiota, we can better understand and potentially modulate the composition of these microbial communities and their interactions with the plant host.

## METHODS

Please see Supporting Information.

## ACKNOWLEDGMENTS

This work was supported by National Institutes of Health grants R35 GM147018 (to A.W.P.) and R35 GM150722 (to T.L.K). Work in the T.L.K. lab is also supported by NSF 774 2422727 and USDA-NIFA 10074268. M.A. was supported by NIGMS T32 GM007464. We thank T.J. Erb (MPI Marburg) for plasmid pTE1887, W.W. Metcalf (University of Illinois at Urbana-Champaign) for *E. coli* WM6026, C.D. Smart (Cornell University) for *Clavibacter michiganensis* ssp. *michiganensis* 0317, H. Crandall (University of Utah) for *Staphylococcus aureus* MN8, and M.P. Miller (University of Utah) for *Saccharomyces cerevisiae* W303 and helpful discussions about yeast genetics. CRAGE plasmids 158207, 158210, 158211 were a gift from Y. Yoshikuni via Addgene. Plasmid pAC-EHER (53262) was a gift from F.X. Cunningham Jr. via Addgene. We thank R.C. Hurrell (University of Utah) for constructing *E. coli* BW25113 Δ*tolC* Δ*bamB*.

## DATA AVAILABILITY

The mass spectrometry data were deposited in the public repository MassIVE via ID MSV000101826.

## AUTHOR CONTRIBUTIONS

The manuscript was written through contributions of all authors. All authors have given approval to the final version of the manuscript.

## TABLE OF CONTENTS GRAPHIC

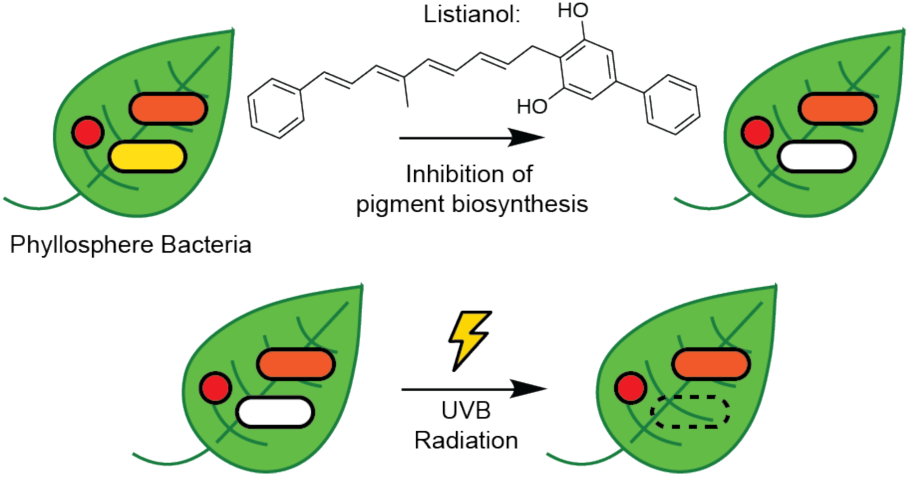

