## Supplementary material for "A Bacterial Natural Product Reshapes Phyllosphere Microbiome Composition by Blocking Carotenoid Biosynthesis": Methods, Figures S1-S19, Tables S2 and S3

### EXPERIMENTAL PROCEDURES

**Routine bacterial culturing**. All strains used in this study are listed in Table S4. *E. coli* strains TOP10 (Thermo Fisher Scientific) and WM6026 were used for plasmid assembly. *Saccharomyces cerevisiae* W303 was used for assembly of BGCs by transformation-associated recombination (TAR). All *E. coli* strains were grown at 37°C on lysogeny broth (LB) medium and supplemented with antibiotics and diaminopimelic acid (DAP) as noted. For *E. coli*, kanamycin and apramycin were used at 50 µg $ml^{-1}$ and chloramphenicol was used at 35 µg $ml^{-1}$. DAP was used at 0.3 mM for *E. coli* WM6026. 50 µg $ml^{-1}$ kanamycin and 100 µg $ml^{-1}$ apramycin were used for *Methylorubrum extorquens* PA1-derived strains. All PPFM strains were grown on a modified version of ammonium mineral salts (AMS) medium[1,2] containing final concentrations of 4 mM phosphate buffer pH 6.8 and 50 mM methanol at 30°C, unless otherwise noted. This version of AMS contains 0.2 g $L^{-1}$ $MgSO_4 \cdot 7H_2O$, 0.2 g $L^{-1}$ $CaCl_2 \cdot 6H_2O$, 0.5 g $L^{-1}$ $NH_4Cl$, 30 µM $LaCl_3$, and 1X trace elements. 500X trace elements contains: 1.0 g $L^{-1}$ $Na_2$-EDTA, 2.0 g $L^{-1}$ $FeSO_4 \cdot 7H_2O$, 0.8 g $L^{-1}$ $ZnSO_4 \cdot 7H_2O$, 0.03 g $L^{-1}$ $MnCl_2 \cdot 4H_2O$, 0.03 g $L^{-1}$ $H_3BO_3$, 0.2 g $L^{-1}$ $CoCl_2 \cdot 6H_2O$, 0.6 g $L^{-1}$ $CuCl_2 \cdot 2H_2O$, 0.02 g $L^{-1}$ $NiCl_2 \cdot 6H_2O$, and 0.05 g $L^{-1}$ $Na_2MoO \cdot 2H_2O$. All Leaf strains were grown on R2A medium supplemented with methanol to a final concentration of 0.5% (v/v).

**Plasmids**. *General cloning procedures.* All primers and plasmids used in this study are listed in Tables S5 and S6. All DNA fragments used in assembly were amplified with Q5 DNA polymerase (NEB, M0419) using either genomic DNA, plasmid DNA or bacteria biomass as the template. Screening PCRs were performed with One*Taq* DNA Polymerase (NEB, M0480). All plasmids were assembled with NEBuilder HiFi DNA Assembly Mix (NEB, E2621), except for plasmids containing BGCs, which were assembled using yeast TAR. *E. coli* WM6026 was used for assembling and maintaining plasmids with the R6K ori and *E. coli* TOP10 was used for all other plasmids. *E. coli* transformants were screened for correct insert sizes using either insert-specific or backbone-specific primers. Correct colonies were used to inoculate overnight cultures grown at 37°C in LB medium. The plasmids were subsequently isolated (Thermo Scientific, K0503) and sequenced using whole-plasmid sequencing (Plasmidsaurus).

*Cloning of the lux operon reporter plasmids*. First, the promoter sequences and the pW34 backbone containing the *luxCDABE* operon, excluding the T7 promoter sequence, were amplified. These fragments were assembled using NEBuilder HiFi DNA assembly kit and transformed into *E. coli* WM6026. Colonies were screened by PCR and sequence-verified.

*Cloning of BGC plasmids by yeast TAR.* All BGC plasmids were designed to incorporate $P_{L/O4/A1}$, an artificial ribosomal binding site, the BGC of interest, and the T7 terminator region. First, we assembled plasmid pAWP865. This plasmid contains $P_{L/O4/A1}$ and the T7 terminator region from pW34 on the pW5Y-Apr backbone for assembly of BGCs using yeast TAR. For each BGC, an artificial ribosomal binding site was designed using the RBS calculator[3], with a target translation initiation rate of 20,000. This RBS was incorporated via overlapping primer tails between the first BGC fragment and the backbone. BGCs were amplified as 3-5 kb fragments with 50-100 bp overlaps between adjacent BGC fragments or backbone fragments. pAWP865 was amplified as 3 backbone pieces.

For assembling BGCs, pAWP865 fragments and BGC fragments were combined at 50 ng $kb^{-1}$, transformed into *S. cerevisiae* W303 using the PEG-LiAc method and plated onto yeast synthetic drop-out medium without uracil (Sigma-Aldrich, Y1501) containing yeast nitrogen base without amino acids and 2% (w/v) glucose. After 3 days, ~50 colonies were resuspended in 30 µL lysis buffer (Thermo Scientific, K0503) containing ~5 0.5mm glass beads. This mixture was vortexed for 1 minute and 2 µL of the mixture was transformed into *E. coli* TOP10. The resulting colonies were screened using BGC-specific primers and plasmids from positive colonies were sequence-verified.

**Identification of BGCs from PPFMs using antiSMASH**. First, we downloaded the genomes of strains within our own strain collection and the *At*-LSPHERE collection from the genera *Methylorubrum* and *Methylobacterium* via the Joint Genome Institute's Integrated Microbial Genomes and Microbiomes (IMG/M) system[4]. These genomes, ~65, were then analyzed using antiSMASH version 6.1.1 to identify putative BGCs. BGCs were analyzed manually to find suitable candidates for heterologous expression, prioritizing BGCs with co-directionality, unknown products, and sizes under 60 kb. We then used the Enzyme Function Initiative-Genome Neighborhood Tool (EFI-GNT)[5] to compare genome neighborhoods of strains that contained our prioritized BGCs to infer possible BGC boundaries for cloning. The gene IDs of the BGCs expressed in this study can be found in Table S1.

**Integration of landing pad into *Methylorubrum extorquens* PA1**. We used conjugation to transform pW17 into *Methylorubrum extorquens* PA1 strain AWP227. *E. coli* EcS17 λpir harboring pW17 was spread on LB medium containing 50 µg ml$^{-1}$ kanamycin and grown overnight at 37°C. Simultaneously, one inoculation loop of AWP227 was spread onto AMS media containing 50 mM methanol and 10% (v/v) nutrient broth, and incubated overnight at 30°C. The following day, one inoculation loop of the *E. coli* was transferred to the plate with AWP227, the cells were mixed until a homogenous lawn was formed, and the plate was incubated overnight at 30°C. The next day, the cell mixture was resuspended in 1 mL AMS medium, and dilutions were plated to AMS medium containing 50 mM methanol and 50 µg ml$^{-1}$ kanamycin. After ~5 days, colonies were patched to AMS medium containing 50 mM methanol and 50 µg ml$^{-1}$ kanamycin to verify resistance to kanamycin. After 2 days, these patches were screened by PCR for insertion of the landing pad. We also screened for pW17 at 2 different locations in the backbone to verify that it was not replicating in AWP227.

**Identification of LP insertion site**. To identify the landing pad insert locations, we used an arbitrary PCR method adapted from Wozniak et al.[6]. In the first round of amplification, an LP-specific primer and a primer containing a random tetramer sequence with a tail (oAWP 2466 and oAWP2403) were mixed with biomass from PA1-LP in a 20 µL OneTaq (NEB) reaction. DNA was amplified for 30 cycles with 30s at 94°C, 1 min at 38°C, and 2 min at 68°C. 0.5 µL of this reaction was used as the template for a second 20 µL OneTaq reaction, which contained a downstream primer in the LP and a primer whose sequence matched the tail of the random tetramer primer (oAWP2467 and oAWP2135). DNA was amplified for 30 cycles with 30s at 94°C, 1 min at 56°C, and 2 min at 68°C. The PCR products were column-purified and sanger sequenced using the downstream LP primer. Final LP insert sites were verified by long-read amplicon sequencing (Plasmidsaurus) of PCR products that spanned the genome regions upstream and downstream of the LP.

**Integration of BGCs into *Methylorubrum extorquens* PA1-LP**. Plasmids containing BGCs were transformed into PA1-LP via conjugation using a triparental mating. The mating was performed as described in the previous section using *E. coli* TOP10 harboring the BGC plasmid, *E. coli* HB101 harboring the mobilization plasmid pRK2013, and PA1-LP as the recipient strain. The resulting mating was plated to AMS medium containing 50 mM methanol and 100 µg ml$^{-1}$ apramycin and after ~5 days, colonies were patched to the same medium. Patches were screened by colony PCR using BGC-specific primers to verify integration of the BGC. We verified that the BGC plasmids were not replicating in PA1-LP by colony PCR at two locations on the pW5Y-Apr backbone.

**Bioluminescence assay**. For each PA1-*lux* reporter strain, ~20 colonies were used to inoculate 6 mL of AMS medium, which was grown overnight at 30°C. Subsequently, cultures were diluted to OD ~0.1 and 1.2 mL aliquots were transferred to deep 96-well plates (Evergreen Scientific). Isopropyl β-d-1-thiogalactopyranoside (IPTG) was added to the wells to final concentrations from 0-2 mM. There were 3 replicates per condition. The plate was covered with a breathable membrane (AeraSeal, Excel Scientific) and incubated at 30°C shaking at 200 rpm for 2 days. Subsequently, 150 µL of each well was transferred to a white 96-well plate (Evergreen Scientific). Luminescence was measured on a SpectraMax i3x plate reader and the results were visualized using Graphpad Prism version 8.0.2.

**Screening of PA1-BGC strains for new metabolites**. ~20 colonies from plates of PA1-LP and PA1-BGC strains grown on AMS medium were used to inoculate 6 mL of AMS medium. These cultures were grown in a shaking incubator at 30°C and 200 rpm overnight. 600 µL of these cultures were used to inoculate 6 mL AMS medium, resulting in an $OD_{600}$ of ~0.1, and these cultures were again grown overnight. The following morning, 600 uL of these subcultures were used to inoculate fresh 6 mL AMS medium. After ~6 hours, expression of the BGCs was induced with the addition of isopropyl β-d-1-thiogalactopyranoside (IPTG) to a final concentration of 1 mM. These cultures were incubated for 4 days at 30°C and 200 rpm before extraction. After 4 days of growth, the cultures were centrifuged to produce cell pellets and cell-free supernatants. Cell-free supernatants were extracted twice with 6 mL of ethyl acetate containing 0.1% (v/v) acetic acid. The cell pellets were resuspended in 1 mL of methanol and sonicated for 15 minutes at 30°C, followed by centrifugation at 21,000 x g for 10 minutes. Both ethyl acetate extracts and the methanol extract were combined and evaporated under nitrogen. Dried extract was resuspended in 200 µL 50% acetonitrile/$H_2O$ before being analyzed by LC-HRMS/MS.

**High Resolution Tandem Mass Spectrometry (LC-HRMS/MS).** Mass spectrometry data were collected using an Agilent Revident Q-TOF coupled to an Agilent Infinity III HPLC system with an Acquity UPLC HSS T3 C18 column (1.8 µm, 2.1 x 50 mm). Solvent A: Water + 0.1 % (v/v) formic acid, Solvent B: Acetonitrile + 0.1% (v/v) formic acid. The sample was eluted from the column using a 10-minute linear solvent gradient: 0-0.1 min, 1% B; 0.1 - 10 min, 1-100% B, followed by 2 minutes of 100% B. The solvent flow rate was 0.4 mL $min^{-1}$ and the injection volume was 10 µL. Mass spectra were collected in positive and negative ion mode in separate injections, using the Auto-$MS^2$ acquisition mode. An MS range of *m/z* 100-1500 and an $MS^2$ range of *m/z* 50-1500, both at 5 spectra/s, were used with the isolation width set to medium (~ *m/z* 4). The collision energy gradient was set to automatic according to *m/z* values of precursor ions. Under the collision energy section, Formula was used with two lines, charge set to “All”, “Slope” is 2.6 and “Offset” is 14.75, and charge set to “All”, “Slope” set to 3.9 and “Offset” to 22.13. The maximum precursors per cycle was set to 5, with the Absolute Precursor Threshold set to 15000 (Relative Threshold 0.015%) and Active Exclusion enabled.

**Large-scale production and purification of listianol from *Methylorubrum extorquens* PA1-Lst**. Two 6 mL exponentially-growing cultures of PA1-Lst were used to inoculate two flasks with 300 mL of AMS medium containing 50 mM methanol. After growing overnight, 100 mL aliquots from these cultures were used to inoculate six 2.8 L baffled flasks containing AMS medium with 50 mM methanol. These cultures were grown for ~6 hours at 30°C and 200 rpm before being induced with the addition of IPTG to 1 mM. After 3 days of additional growth, the cultures were centrifuged and the supernatant discarded. Listianol is primarily found associated with the cell pellet after centrifugation, likely due to its non-polar nature. The pellets were resuspended in water and lyophilized. The lyophilized pellets were resuspended in 30 mL of methanol, sonicated for 10 minutes, and centrifuged. Methanol extracts were combined, and this process was repeated 4-5 times, until there was no detectable absorbance at 350 nm in the methanol post-sonication. The combined methanol extracts were concentrated *in vacuo* to produce a yellow oil. This oil was resuspended in 5 mL of 40% acetonitrile/$H_2O$ and loaded onto a Sep-Pak 5g C18-SPE cartridge (Waters). The column was then washed with ~50 mL of 40% acetonitrile/$H_2O$. The listianol-containing fraction was then eluted from the column with 100% acetonitrile. This eluate was dried, resuspended in 100% acetonitrile, and separated using an Agilent 1260 Infinity liquid chromatography system equipped with a Waters XBridge Phenyl column (10mm x 100mm, 5 µm particle size). Listianol was purified using two separate isocratic chromatography methods. The first method, 4 mL $min^{-1}$ flow rate, 0-10 min isocratic at 60% acetonitrile/40% $H_2O$, 10-10.5 min gradient 60-100% acetonitrile, 10.5-16 min isocratic 100% acetonitrile, yielded impure fractions of listianol. These impure fractions were pooled, concentrated *in vacuo*, resuspended in methanol, and separated by a second isocratic method: 5 mL $min^{-1}$ flow rate, 0-15 min isocratic 72% MeOH and 28% $H_2O$, 15-15.5 min gradient 72-100% MeOH, 15-18 min isocratic 100% MeOH. Pure fractions were pooled, concentrated *in vacuo*, and lyophilized to yield listianol as a white solid (2.5 mg).

**Listianol chemical characteristics**. $C_{28}H_{26}O_2$. White powder. UV/Vis: $\lambda_{max}$ 260 nm, 333 nm, 349 nm, 366 nm. High-resolution MS: $[M – H]^-$ calc. 393.1860, observed 393.1867, +1.78 Δppm. NMR data in DMSO-$_{d6}$: Figures S5-8, Table S2**.**

**NMR experiments**. Purified listianol was dissolved in DMSO-$d_6$ and transferred to a 3mm NMR tube. Initial NMR experiments, $^1H$, $^{13}C$-$^1H$ gCOSY, $^{13}C$-$^1H$ HSQC, and $^{13}C$-$^1H$ HMBC, were performed on an Agilent DirectDrive 500 with a high-sensitivity HCN cold probe. Further $^1H$ and $^{13}C$-$^1H$ gCOSY NMR experiments were conducted on a Varian VNMRS 800 for increased resolution in the aromatic region. All NMR data were analyzed using MestReNova software.

**Inverse stable isotopic labeling (InverSIL) experiments**. InverSIL experiments with *Methylorubrum extorquens* PA1-derived strains were performed as previously described[55]. Exponentially-growing cultures grown on AMS medium with 50 mM unlabeled methanol were centrifuged and the pellets washed twice with AMS medium containing no carbon source. After washing, the pellets were resuspended in fresh AMS medium containing no carbon source and used to inoculate cultures to an OD of 0.05. To these cultures, unlabeled methanol or $^{13}C$-methanol were added to a final concentration of 50 mM. When indicated, unlabeled precursors were added to a final concentration of 100 µM. Cultures were then grown for 3 days at 30°C and 200 rpm. After 3 days, the cultures were extracted with ethyl acetate and methanol, then analyzed via LC-HRMS/MS. All InverSIL experiments were performed 3 or more times.

**Quantification of carotenoid biosynthesis inhibition by listianol**. For all experiments, strains were grown in R2A-MeOH medium at 30°C, except for *Staphylococcus aureus* MN8, which was grown in tryptic soy broth (TSB) medium at 37°C. Exponential-phase cultures were used to inoculate 18 mm culture tubes containing 8 mL media and listianol at the tested concentrations, in duplicate. *S. aureus* cultures were inoculated with 8 µL of exponential phase cultures and other strains were inoculated with 80 µL. PPFMs were grown for 2 days and non-PPFM strains were grown for ~20 hours. After growth, cultures were pelleted and the cell pellets were washed once with phosphate buffered saline (PBS), pH 7.4 to remove excess media. Cell pellets were then resuspended in 1 mL of MeOH at 55°C and sonicated for 15 minutes at 55°C. These extracts were centrifuged, and the supernatants were transferred to a glass test tube. This process was repeated twice more for all cultures to remove all pigments from the cell pellets. Supernatants were combined, dried under nitrogen, and resuspended in 200 µL of MeOH. 150 µL of these resuspensions were transferred to black 96-well plates (Evergreen Scientific) and absorbance was measured on a SpectraMax i3x plate reader. An absorbance wavelength of 495 nm was used for PPFMs and 445 nm for non-PPFMs, coinciding with approximate maximum absorbance wavelengths for carotenoids produced by the tested strains. The data was analyzed using Graphpad Prism version 8.0.2 to generate dose-response curves and to calculate $IC_{50}$ values. Two independent experiments were performed.

**Analysis of listianol inhibition of carotenoid biosynthesis enzymes heterologously expressed in *E. coli***. Exponential-phase cultures of *E. coli* expressing carotenoid biosynthesis genes were diluted 1000-fold into duplicate 8 mL cultures of LB containing the appropriate antibiotic. Listianol, or a methanol vehicle control, was added to these cultures at a volume of 12 µL to the indicated final concentration and the cultures were grown overnight at 37°C and 200 rpm. These cultures were pelleted after ~20 hours of growth, due to delayed pigment development of the *E. coli* strains expressing CrtB and CrtI. Once pelleted, the pigments were extracted as described in the previous section, except that 5 rounds of extraction were needed to extract all the pigments from the *E. coli* strain expressing CrtB and CrtI. The methanol extracts were combined, dried under nitrogen, resuspended in 200 µL of methanol and analyzed via LC-HRMS/MS. These extracts were separated using a different liquid chromatography method than other extracts: Solvent A: Water + 0.1 % (v/v) formic acid, Solvent B: Acetonitrile + 0.1% (v/v) formic acid, Solvent C: Isopropanol + 0.1% (v/v) formic acid. The sample was eluted from the column using the following solvent gradient: 0-0.1 min, 60% A, 20% B, 20% C; 0.1 - 10 min, gradient to 0% A, 50% B, 50% C; 10-20 min, isocratic 0% A, 50% B, 50% C. The solvent flow rate was 0.3 mL $min^{-1}$ and the injection volume was 10 µL. The HPLC instrument and column were the same as in previous experiments. The products of these strains were quantified by first extracting the UV-Vis data at their corresponding maximum absorbance wavelength: 288 nm for CrtM and CrtB products, 440 nm for CrtMN products and 465 nm for CrtBI products. Then, area under the curve was calculated for each product peak. These data were plotted in Graphpad Prism version 8.0.2. Two independent experiments were performed.

**Quantification of growth inhibition by listianol**. The broth microdilution method was used to determine the minimum inhibitory concentration of listianol against tested strains. The media, temperature and starting OD for each strain tested can be found in Table S7. All stains were grown in cation-adjusted Mueller-Hinton broth (Sigma-Aldrich), unless they grew poorly in that medium, in which case, they were grown in R2A medium. Inoculum for the assay came from exponentially growing cultures. 150 µL of the tested strains were added to the interior wells of a 96-well plate. Listianol was added to a final concentration of 200 µM to one row of wells and wells were serially diluted 2-fold. Strains were tested in triplicate at each concentration. The cultures were grown overnight shaking in a SpectraMax i3x plate reader. OD readings were taken every 16 minutes for 18 hours. Data were plotted and analyzed in Graphpad Prism 8.0.2 to calculate $IC_{50}$ values. Two independent replicates were performed for each strain.

**UV susceptibility testing.** Exponential-phase cultures were used to inoculate cultures of the tested strains in R2A-MeOH media. Listianol was added to a final concentration of 1 µM and the cultures were incubated overnight at 30°C. Subsequently, cultures were pelleted and the pellets were washed twice with PBS. Cell pellets were resuspended in PBS to ~1 x $10^7$ CFU $mL^{-1}$. 1 mL aliquots were transferred a clear polystyrene 24-well plate (Celltreat) and exposed to narrowband UVB radiation for 6 hours with the lids on. Non-PPFMs were irradiated with 1.6 W $m^{-2}$ and PPFMs were irradiated with 3.6 W $m^{-2}$. The UVB radiation was provided by a 36 W lamp (CureUV, 202226) equipped with a narrowband UVB bulb (Phillips, TUV PL-L 36W/4P). Irradiance was measured with a UV light meter (De-Power, AH-UVCBA) placed under a 24-well plate polystyrene lid. UV intensity was altered by changing the distance of the plate from the lamp. As a negative control for UV treatment, an additional plate was prepared as described but was covered in aluminum foil during the UV treatment. After irradiation, samples were serially diluted and 5 µL was spotted on R2A + MeOH medium in triplicate. Plates were incubated at 30°C for 4 days, then colonies were counted. Percent survivability was calculated by dividing CFUs of UV-treated cells by CFUs of cells from the plate covered in aluminum foil. Data were analyzed using Graphpad Prism 8.0.2. Two independent replicates were performed for each strain.

**Plant growth conditions**. Wild-type seeds of the natural accession of *Arabidopsis thaliana* Ey1.5-2 were surface sterilized using a 70% EtOH and 0.05% Triton mixture for 5 minutes. Seeds were washed with 100% EtOH and left to air dry overnight. Seeds were then spread across ½ MS Agar (PhytoTech Labs, M5531) in square plates (Fisher Scientific, FB0875711A). Seeds were then stratified for 7 days in the dark at 4 °C. After stratification, seeds were grown under short-day conditions (8 hours light, 16 hours dark) in an AR41L3 Percival chamber (Percival Scientific) with a PAR (Photosynthetic Active Radiation) of 145 µmol $m^{-2}$ $s^{-1}$ and temperature of 23°C. Six-day-old seedlings were then transplanted into individual wells of a 24-well plate (Greiner Bio-One, 6621665). Seventeen-day-old plants (four-days post inoculation), were moved into an AR-1015L3LED Percival chamber (Percival Scientific) with a PAR of 70-165 µmol $m^{-2}$ $s^{-1}$ and temperature of 23°C. For six days, the plants were exposed to narrowband-UVB radiation at different intensities during the entire 8-hour light period: 0, 0.5 W $m^{-2}$, 1.0 W $m^{-2}$, and 2.2 W $m^{-2}$. UVB radiation was provided by two bulbs (TL40W/01-RS, Phillips) placed above the plants and the irradiance was measured with a UV light meter (De-Power, AH-UVCBA) placed under the 24-well plate polystyrene lids. UV intensity was altered by changing the distance of the plate from the lamp.

**Inoculation of plants with the SynComs.** The protocol for inoculating plants with SynComs was adapted from Emmenegger et al.[7]. SynCom strains were struck out from frozen stocks onto R2A medium supplemented with 0.5% (v/v) methanol and incubated at room temperature, ~22°C, for 5 days. Strains were resuspended in 6 mL 10mM $MgCl_2$ and vortexed for ~1 minute each. Suspensions were diluted to 0.2 $OD_{600}$ and combined in equal volumes to form the SynComs. The SynComs were then diluted 10-fold and the OD adjusted to 0.02. Thirteen-day-old seedlings were inoculated with 200 µL of the SynComs or 200 µL of $MgCl_2$. Inoculum CFUs were determined by plating 50 µL of SynCom dilutions onto R2A medium supplemented with 0.5% (v/v) methanol and incubating the plates at room temperature. Colonies were counted after 7 days.

**Determining bacterial plant colonization by colony-forming units.** The protocol for assessing the SynCom plant colonization was adapted from Emmenegger et al.[7]. After the six days of exposure to UV, the 23-day-old plants were harvested by cutting the phyllosphere portion of the plant and transferring it to a pre-weighed 2 mL tube containing 400 µL of PBS, pH 7.4 and a 5 mm sterile glass bead. Plant weights were recorded and then the plants were homogenized for 45 s at 25 Hz (TissueLyser II, Qiagen), after which an additional 400 µL of PBS was added. The homogenized plant mixture was serially diluted 10-fold in PBS. 50 µL of the $10^{-3}$, $10^{-4}$ and $10^{-5}$ dilutions were plated to R2A agar supplemented with 0.5% methanol. The plates were incubated at room temperature for 7 days, after which CFUs for each strain were counted. When strains were not detected on the dilution plates, their count was recorded as 0.9 at the lowest dilution counted for that plant. For each plant, percent abundance of a SynCom strain was calculated by dividing the CFUs of that strain by the sum of the CFUs for each strain in the SynCom.

For the WT-SynCom, PA1-LP and *Methylobacterium* sp. Leaf456 were indistinguishable. To determine the relative abundance of each, 5 µL of the dilution series was plated onto both AMS media supplemented with methanol and AMS media supplemented with methanol and 50 µg $ml^{-1}$ kanamycin. PA1-LP is able to grow on kanamycin, due to the presence of the landing pad, while Leaf456 is not. Both strains are able to grow in the absence of kanamycin. CFUs were counted on each plate after 7 days and the PA1-LP CFUs were divided by the total PPFM CFUs from the plate without kanamycin to give the relative abundance of each strain. These relative abundances were multiplied by the total PPFM count from the dilution plates on R2A media to yield CFUs for each strain. In one instance, the CFUs from the AMS supplemented with kanamycin plate were higher than the CFUs from the AMS plates without kanamycin, making the relative abundance calculations impossible. In this case, the average relative abundance of each PPFM from replicate plants was used for calculating CFUs.

**Microbial community analysis.** Analyses were conducted in R v4.5.1 using tidyverse packages (dplyr v1.2.1, tidyr v1.3.1, ggplot2 v4.0.2, readr v2.2.0, readxl v1.4.5), with additional support from purrr v1.2.2, broom v1.0.10, and car v3.1-3. Community analyses were performed using vegan v2.7-3, with phyloseq v1.52.0 and DESeq2 v1.48.2 for data structuring.

Principal coordinates analysis (PCoA) was conducted on Bray–Curtis dissimilarities calculated from relative abundance data. For visualization, ordination was restricted to experiment 2 at UV = 2.2 (high UV), and the first two axes were plotted with variance explained derived from eigenvalues.

Taxon-specific responses were analyzed using linear models of the form:

Relative abundance∼UV×Treatment\text{Relative abundance} \sim \text{UV} \times \text{Treatment}Relative abundance∼UV×Treatment

where UV dose was treated as a continuous predictor and treatment (listianol presence vs. absence) as a categorical factor. Interaction terms were included to assess whether treatment modified the response to UV. Model fits were evaluated using standard ANOVA, and *p*-values for interaction terms were extracted and displayed on plots.

**SUPPORTING FIGURES**

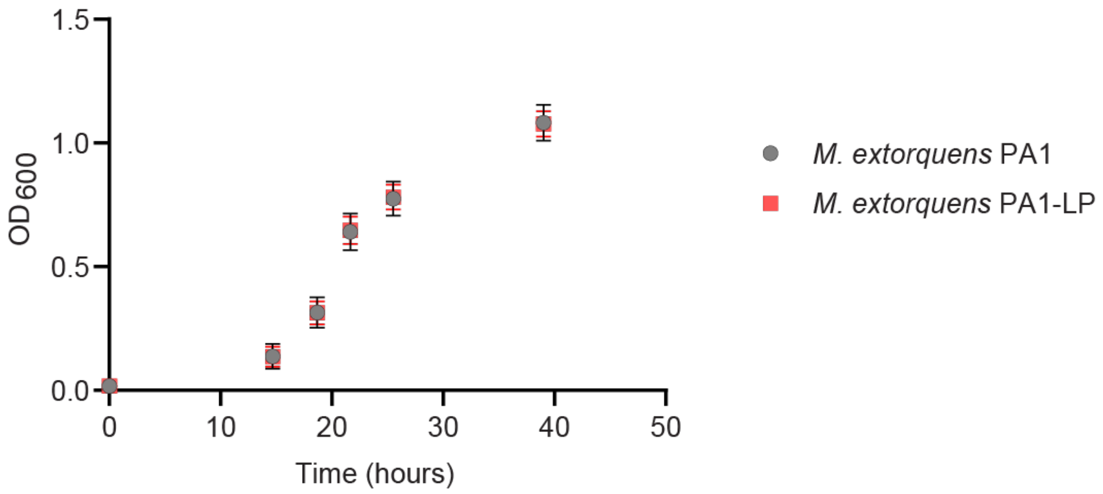

**Figure S1. Comparing the growth of *M. extorquens* PA1 and PA1-LP.** Growth ($OD_{600}$) over a 40 hour period of *M. extorquens* PA1 (grey circles) and *M. extorquens* PA1-LP (red squares). The error bars represent the standard deviation (n=3).

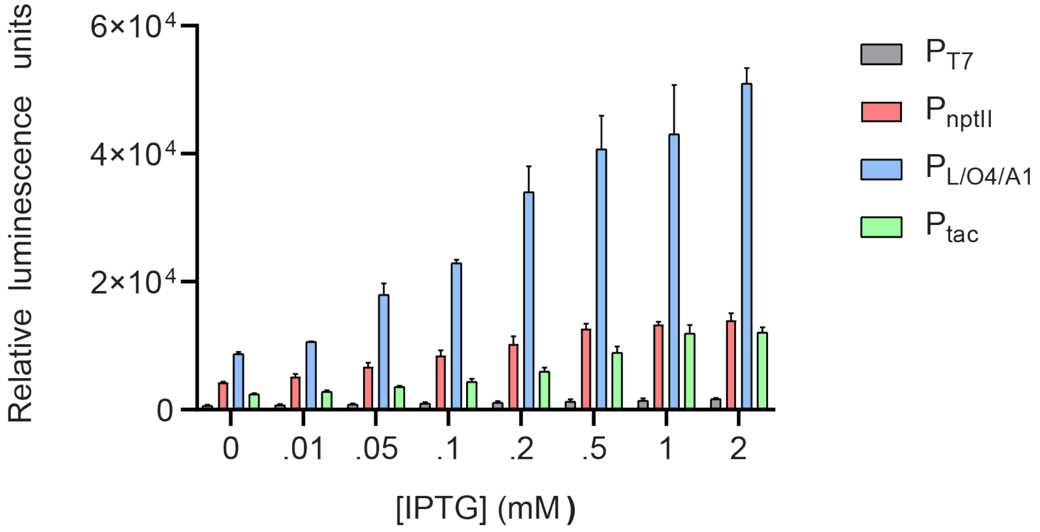

**Figure S2. *lux* operon promoter test in PA1-LP.** Testing four different promoters inserted upstream of the *lux* operon. Relative luminescence after two days of growth in the indicated concentrations of IPTG is shown. The error bars represent the standard deviation (n=3).

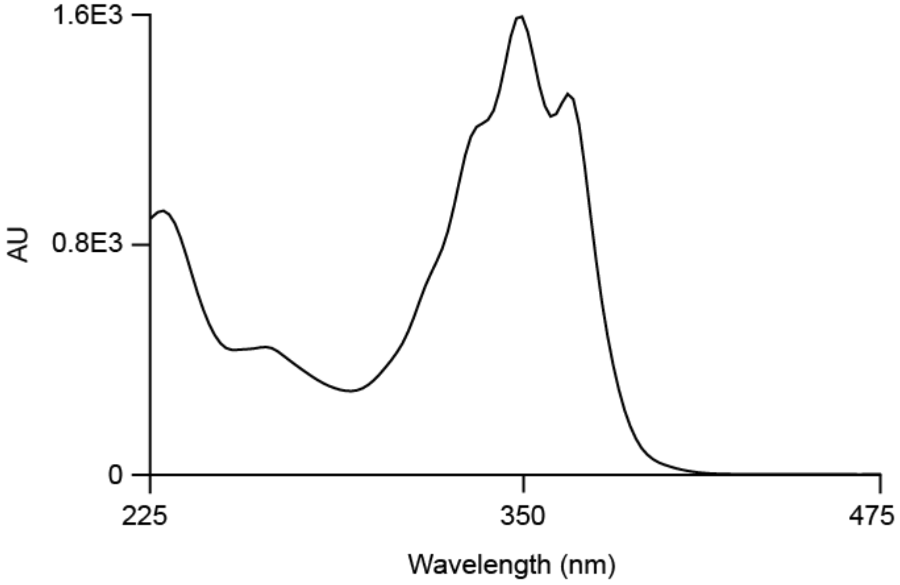

**Figure S3. Absorbance spectrum of listianol.**

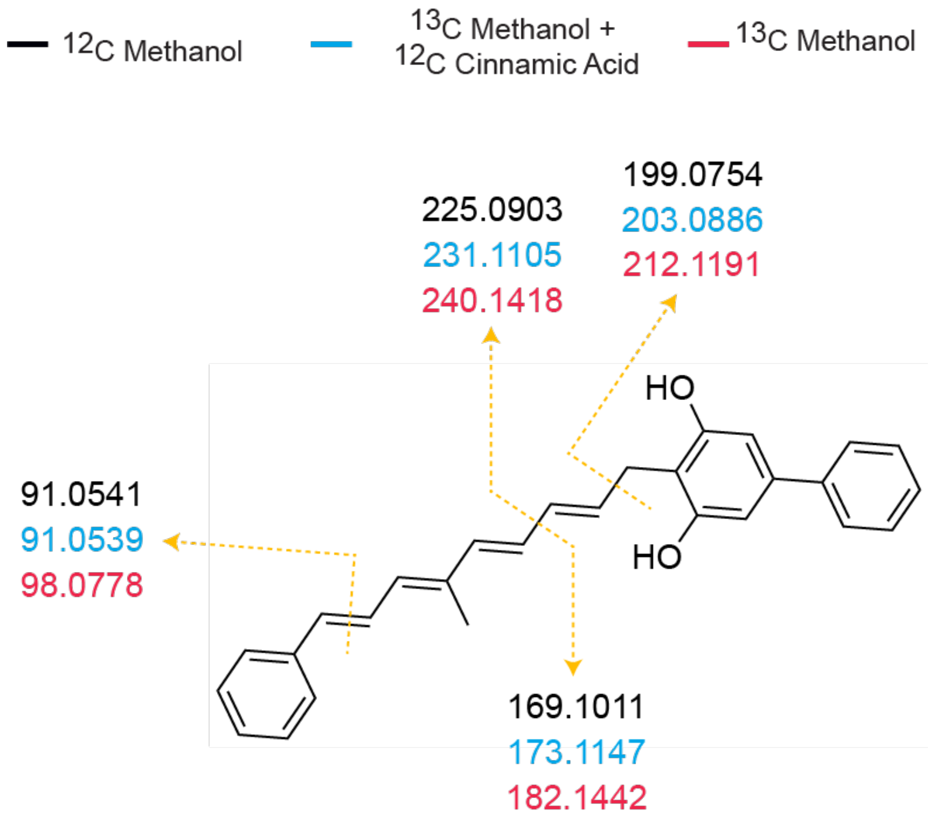

**Figure S4. Key MS$^2$ fragments of listianol.** Shown are MS$^2$ fragments of listianol grown in different InverSIL conditions, where the numbers represent *m/z* values of key fragments. This data was acquired in positive-ion mode and the *m/z* value colors refer to the InverSIL condition the culture was grown in.

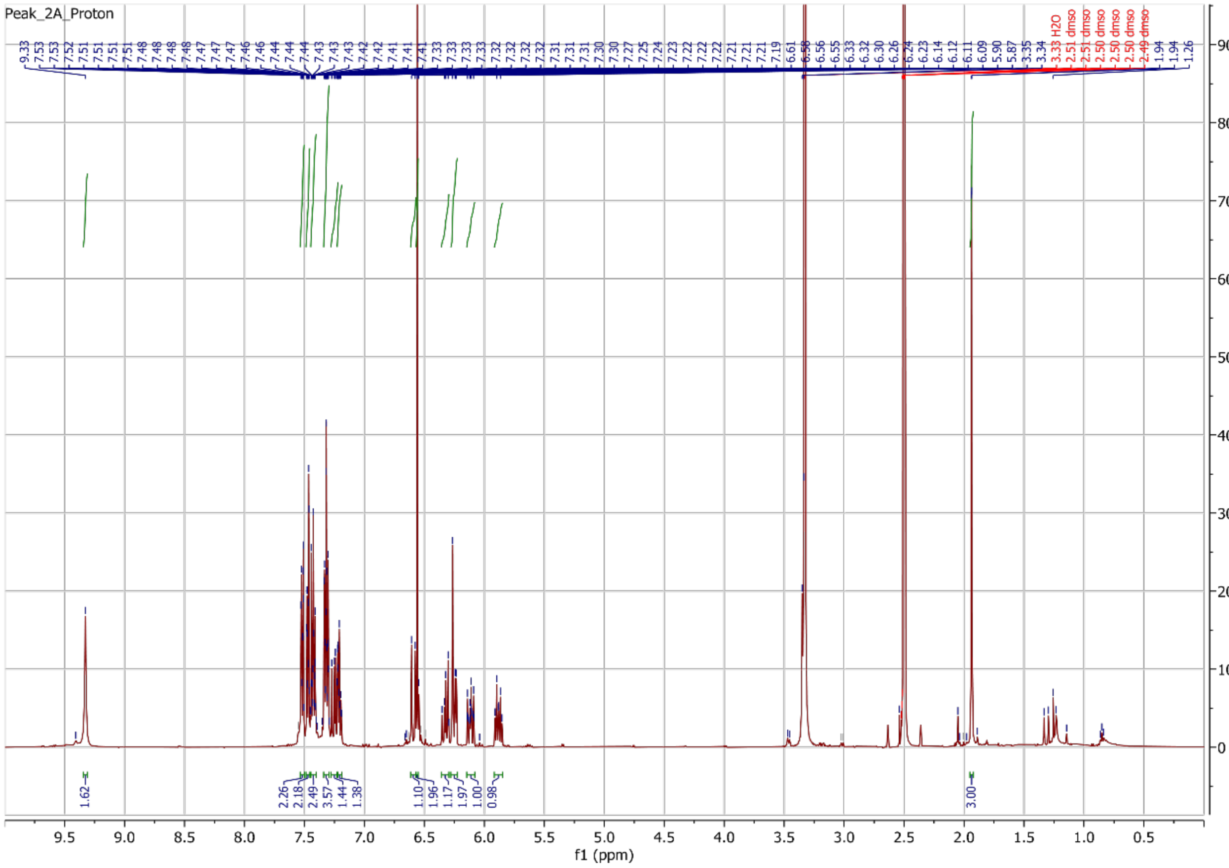

**Figure S5. $^{1}H$ NMR spectrum of listianol in DMSO-$d_6$ (500 MHz).**

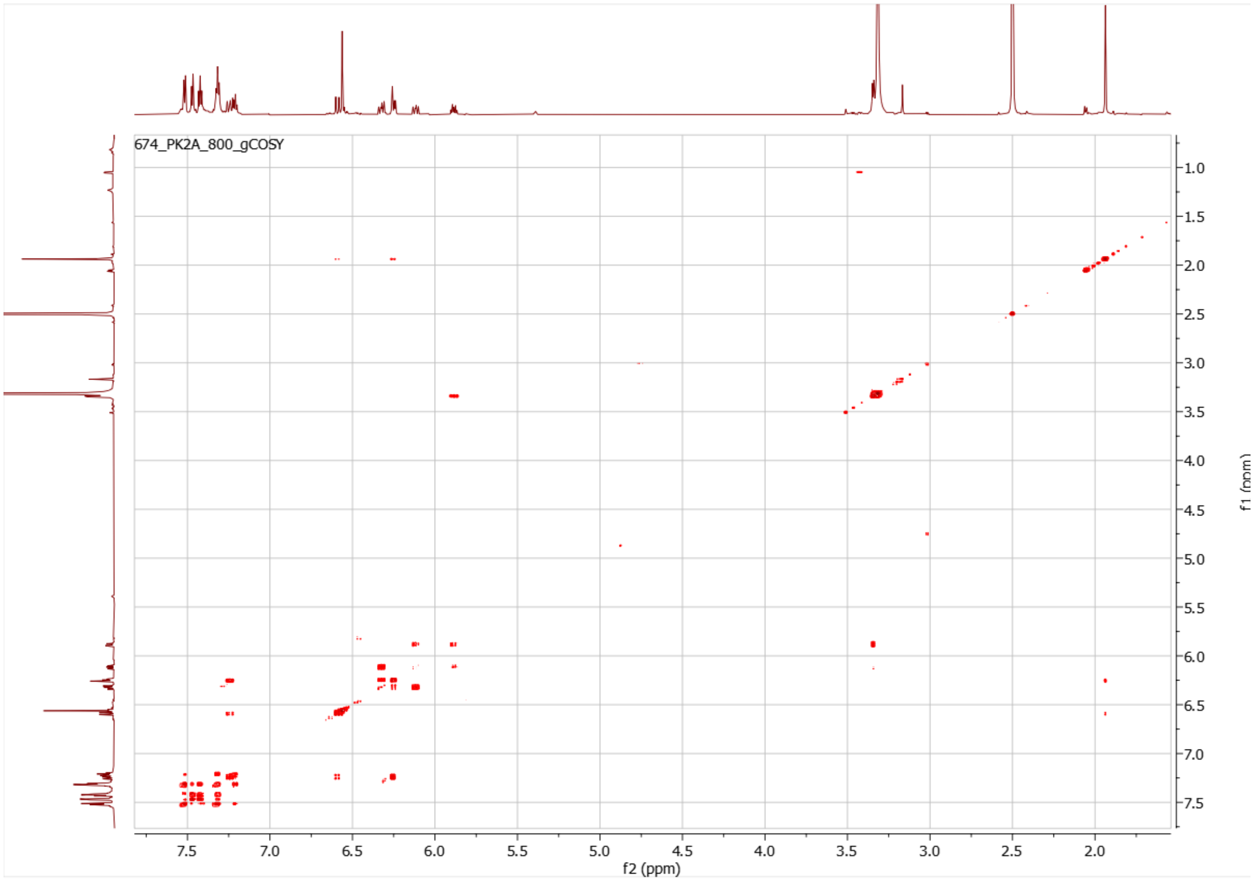

**Figure S6. $^{1}H$-$^{1}H$ gCOSY spectrum of listianol in DMSO-$d_6$ (800 MHz).**

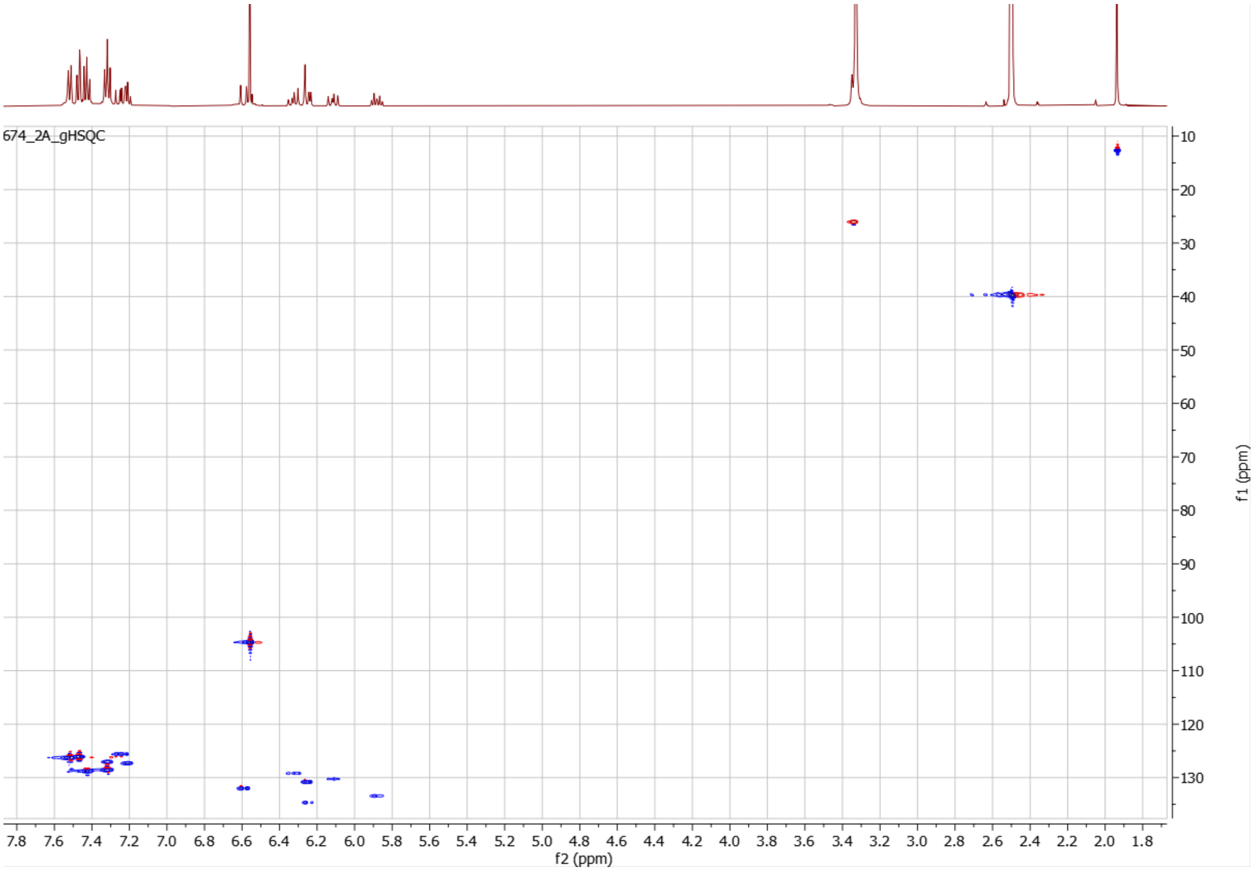

**Figure S7. $^{1}H$-$^{13}C$ gHSQC NMR spectrum of listianol in DMSO-$d_6$ (500 MHz).**

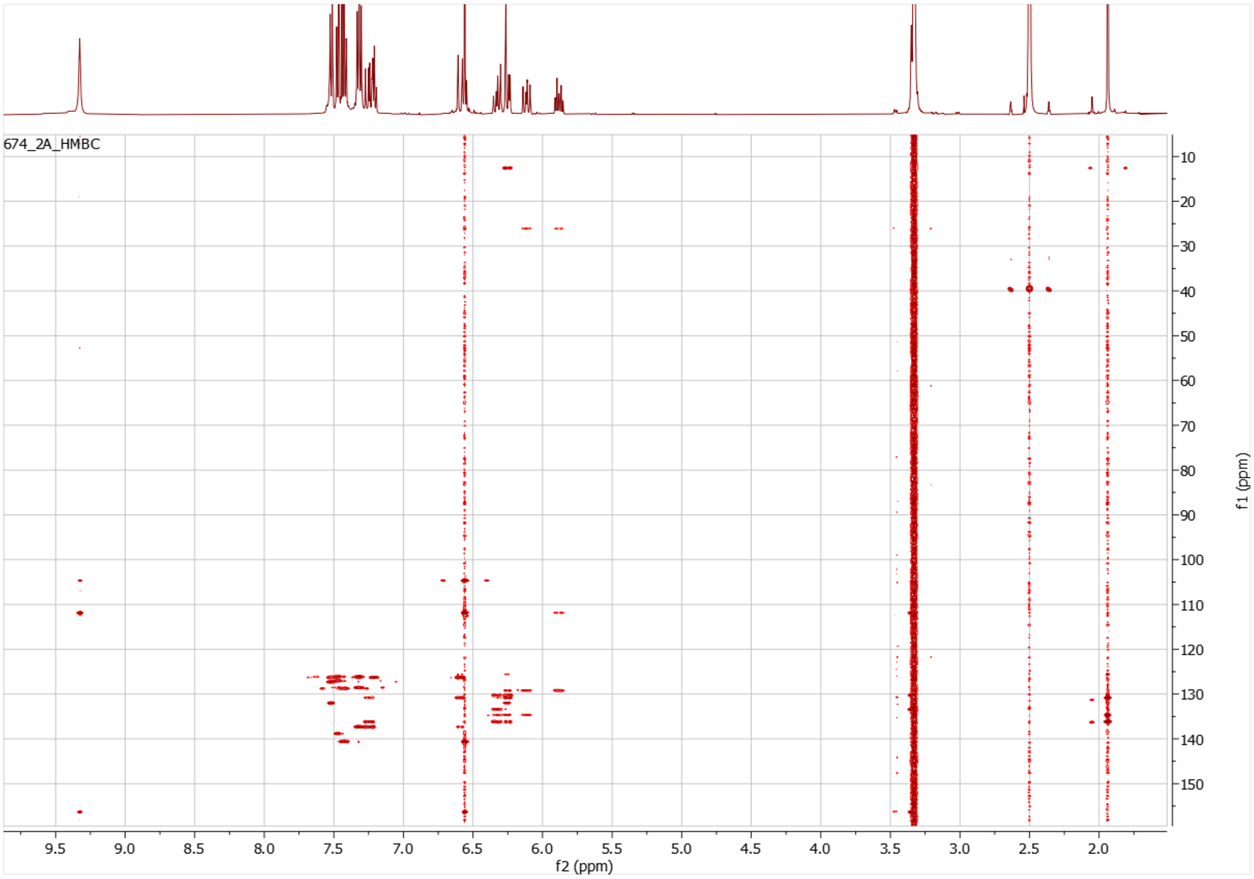

**Figure S8. $^{1}H$-$^{13}C$ HMBC NMR spectrum of listianol in DMSO-$d_6$ (500 MHz).**

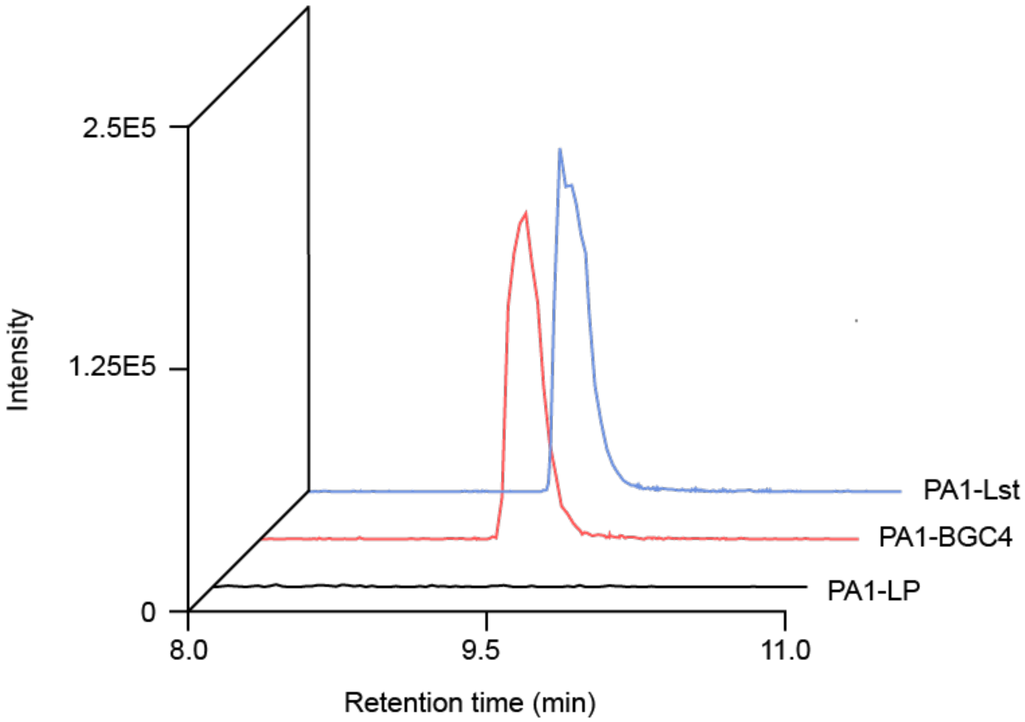

**Figure S9. PA1-BGC4 and PA1-Lst both produce listianol.** Extracted ion chromatogram of listianol, *m/z* of 393.1870 ± 5ppm in negative ion mode from extracts of the indicated cultures.

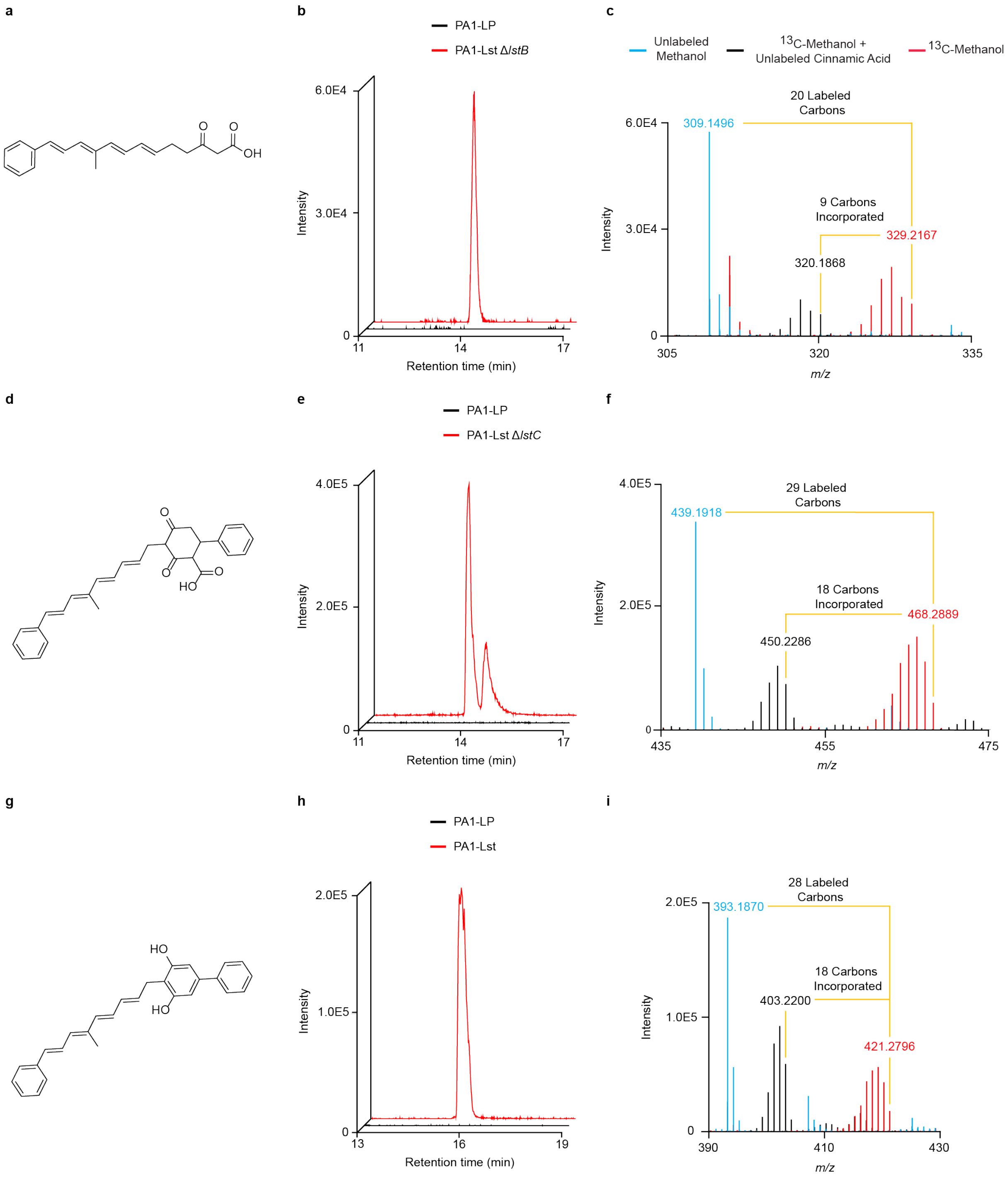

**Figure S10. Supporting evidence for proposed biosynthesis scheme.** Structures of **a**, the predicted PKS product, **d**, the pre-resorcinol carboxy-cyclohexanedione intermediate, and **g**, listianol. Extracted ion chromatograms in negative-ion mode for the predicted *m/z* for the metabolite shown in the same row, **b**, *m/z* 309.1496 ±5 ppm, **e**, 439.1918 ±5 ppm, **h**, 393.1867 ±5 ppm. The two peaks in panel **e** likely arise from tautomerization, as has previously been seen with carboxy-cyclohexanedione intermediates[30]. **c**, **f**, and **i,** Overlaid mass spectra showing the incorporation of cinnamic acid into the metabolite shown in the same row when **c,** PA1-LstΔ*lstB*, **f**, PA1-LstΔ*lstC*, and **I**, PA1-Lst are grown under the conditions indicated in **c**.

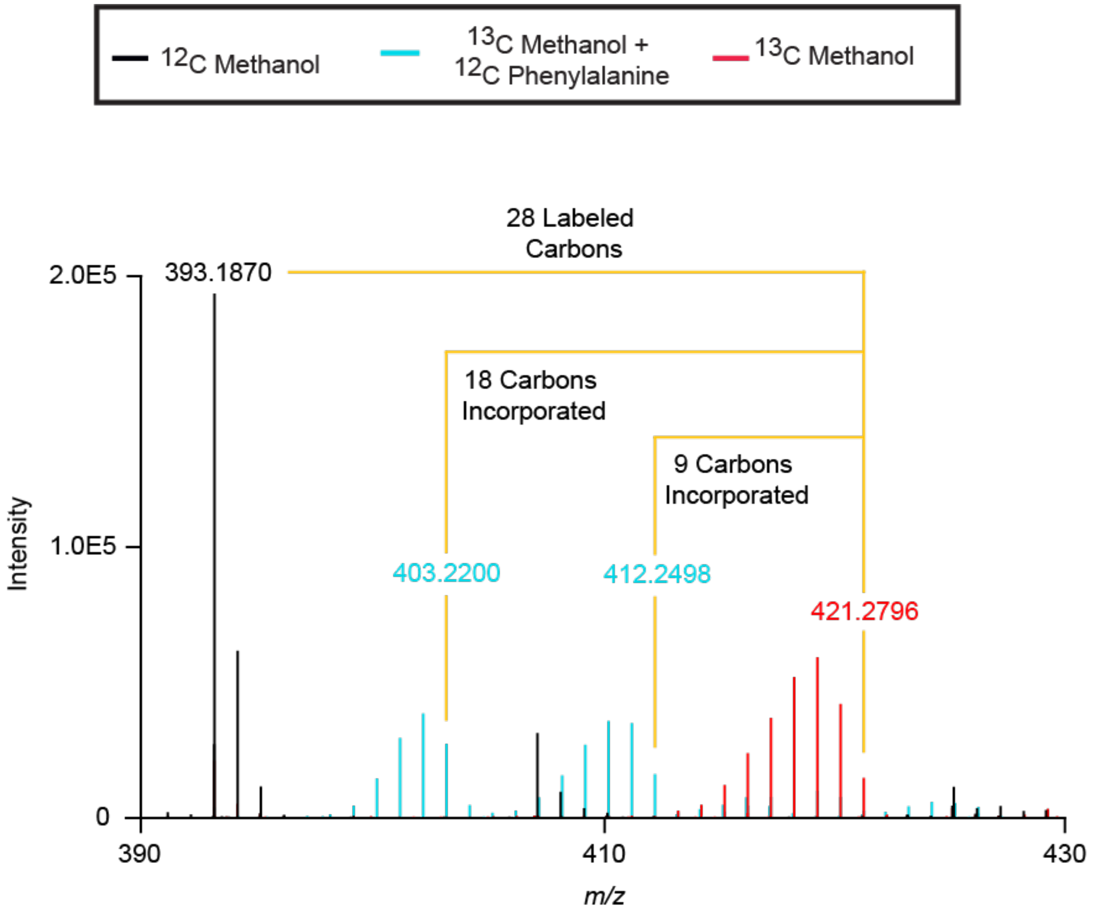

**Figure S11. Listianol incorporates two molecules of phenylalanine.** Overlaid mass spectra showing listianol *m/z* when grown on normal isotopic abundance methanol, $^{13}C$ methanol, and $^{13}C$ methanol supplemented with normal isotopic abundance phenylalanine.

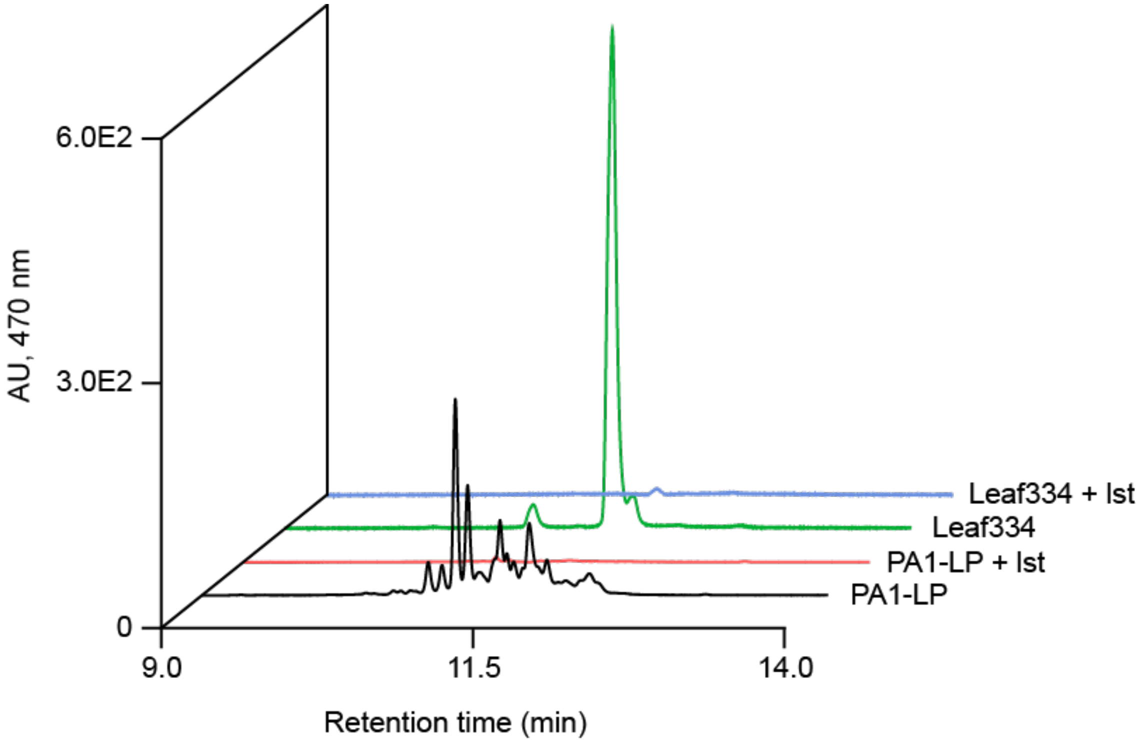

**Figure S12. Listianol treatment eliminates production of carotenoids.** HPLC UV-Vis traces of *M. extorquens* PA1-LP and *Cellulomonas* sp. Leaf334 grown in the absence or presence of 1 μM listianol. Traces were extracted at 470 nm, which is near the maximum absorbance for the carotenoids produced by these strains.

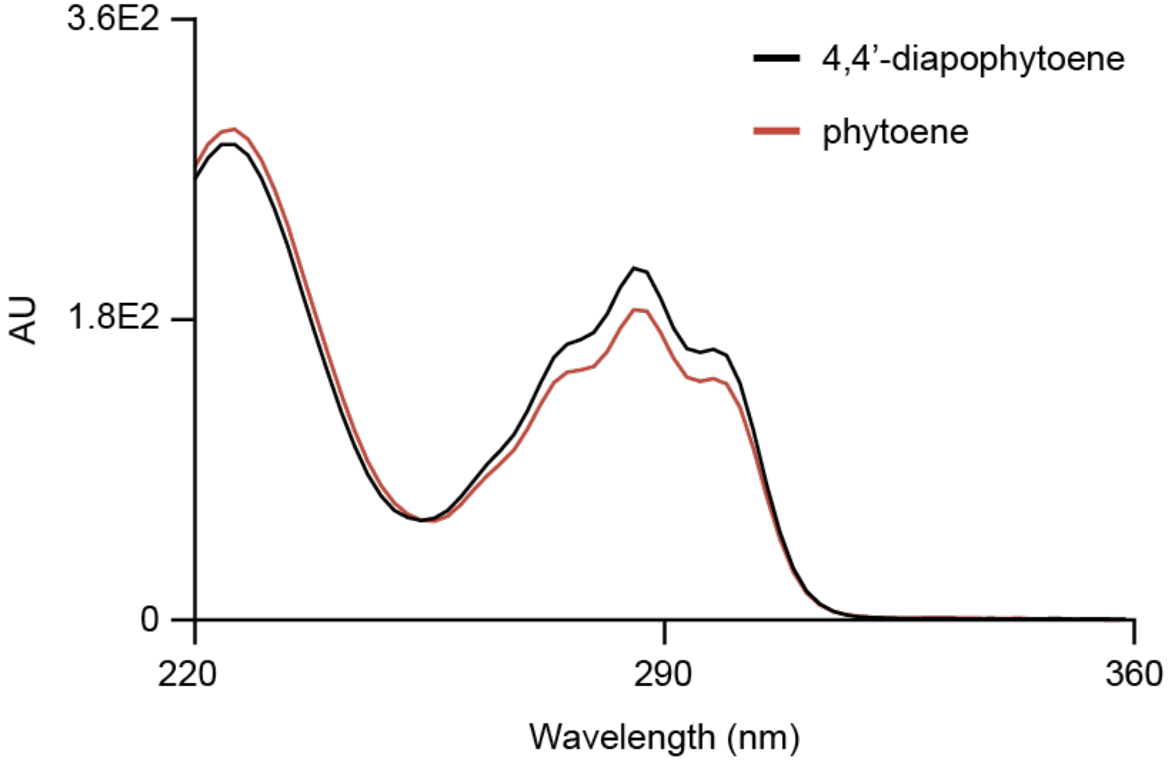

**Figure S13. Absorbance spectra of the products of CrtM and CrtB**. Overlaid UV-Vis spectra of the products of CrtM (4,4'-diapophytoene) and CrtB (phytoene). The y-axis represents absorbance intensity in absorbance units (AU).

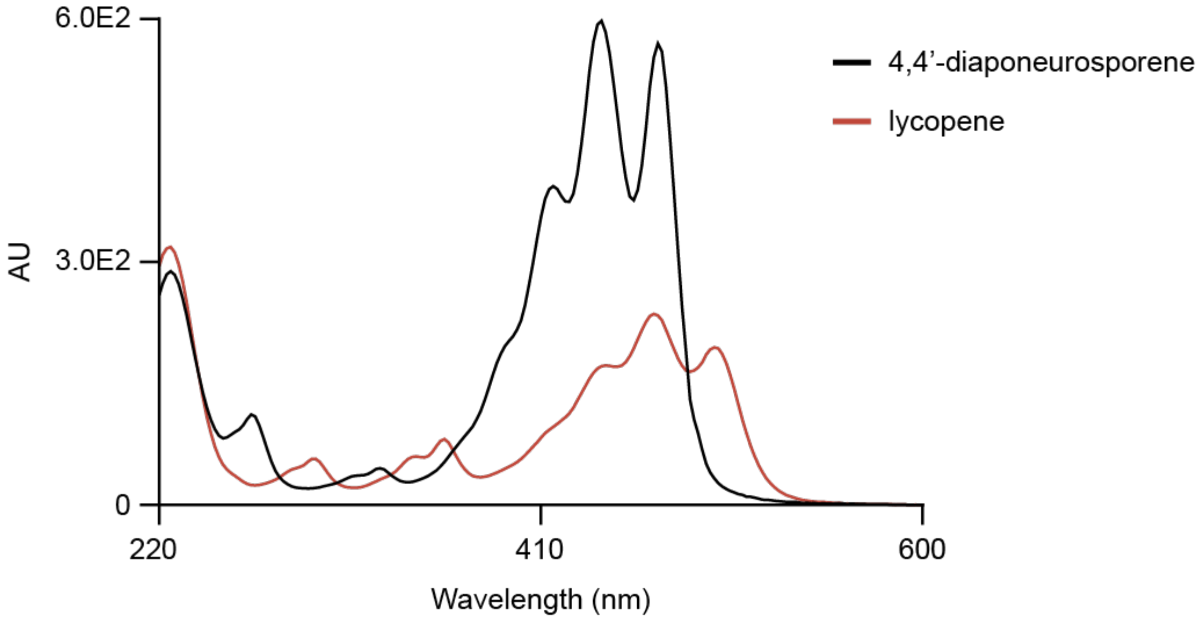

**Figure S14. Absorbance spectra of the products of CrtMN and CrtBI**. Overlaid UV-Vis spectra of the products of CrtMN (4,4’-diaponeurosporene) and CrtBI (lycopene). The y-axis represents absorbance intensity in absorbance units (AU)

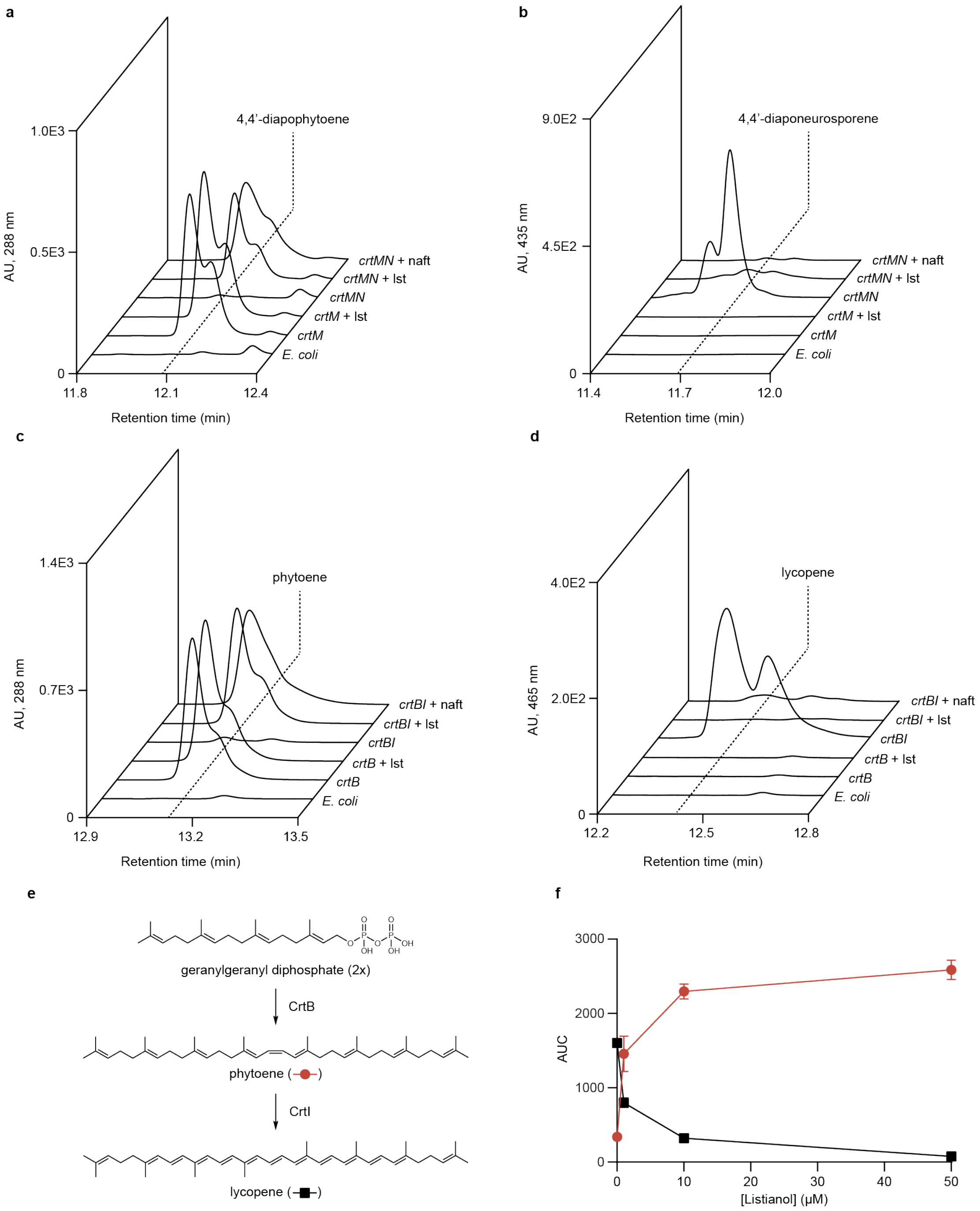

**Figure S15. Evidence for listianol targeting the desaturase enzymes CrtN and CrtI in carotenoid biosynthesis. a-d,** Extracted UV-Vis chromatograms of *E. coli* strains expressing CrtM, CrtMN, CrtB, or CrtBI grown in the absence or presence of 100 μM listianol or 5 μM naftifine. Each enzyme product is shown in a separate panel: **a,** 4,4'-diapophytoene, **b,** 4,4'-diaponeurosporene, **c,** phytoene, and **d,** lycopene. The y-axis represents the absorbance intensity in absorbance units (AU), extracted at the indicated wavelength. **e***,* Scheme illustrating the first steps of C40-carotenoid biosynthesis. **f,** Dose-dependent inhibition of CrtI in *E. coli* – *crtBI* by listianol, showing disappearance of lycopene (black squares) and accumulation of phytoene (red circles). The y-axis represents the area under the curve (AUC) for each product when analyzed by HPLC. CrtB products were extracted at 288nm and the CrtBI products at 465nm.

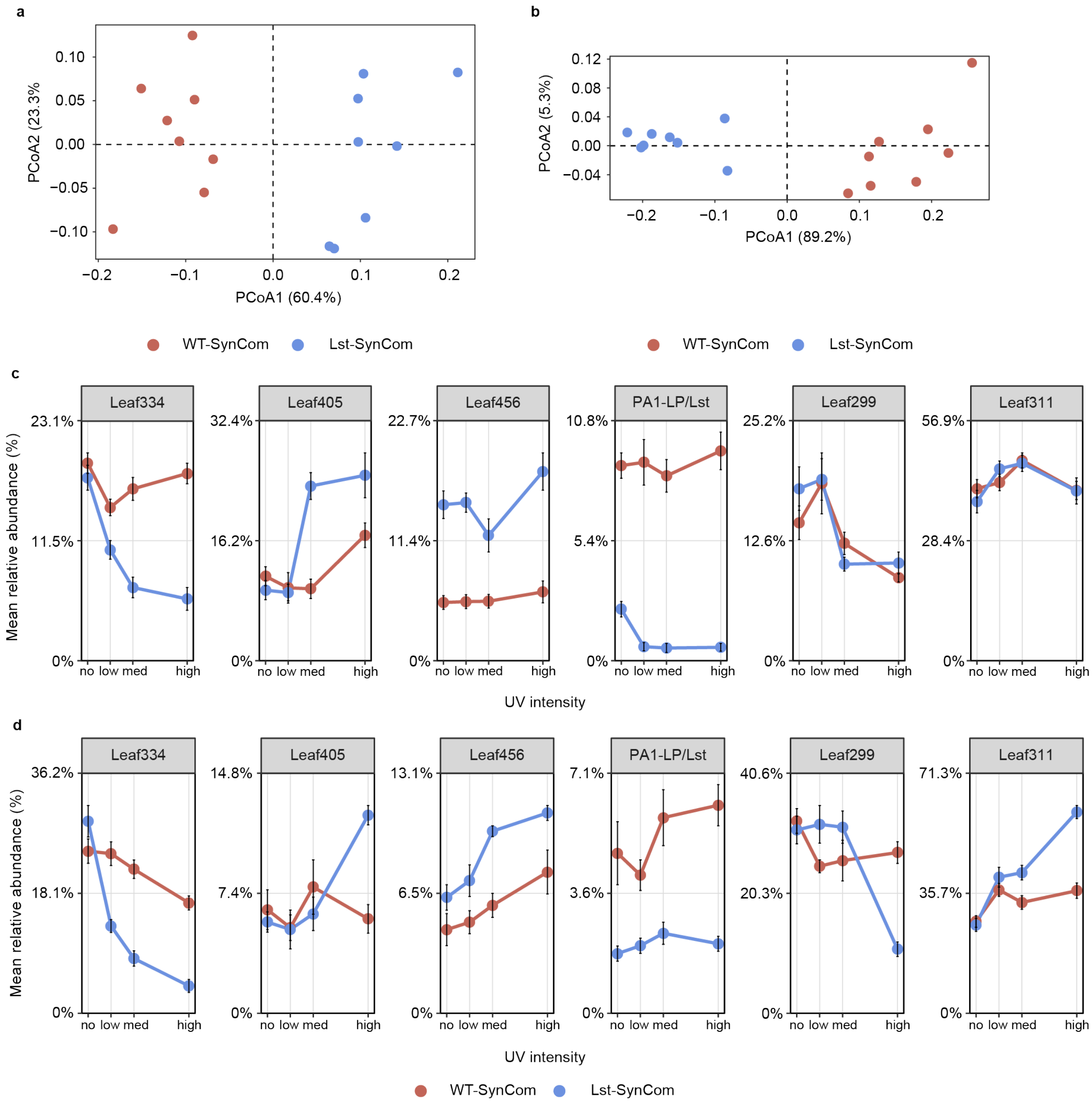

e

| | Leaf334 | Leaf405 | Leaf456 | PA1-LP/Lst | Leaf299 | Leaf311 |
|---|---|---|---|---|---|---|
| *p*-value | <0.001 | 0.00605 | 0.02 | - | 0.00132 | 0.0105 |

**Figure S16. Additional independent replicates of *in planta* UV experiments, supporting the ability of listianol to alter the composition of *A. thaliana* SynComs under UV. a** and **b,** PCoA plots from independent replicate experiments comparing bacterial communities recovered from plants inoculated with WT-SynCom and Lst-SynCom and grown under high-UV irradiation. **c** and **d,** Mean relative abundances for each strain in either the WT- or Lst-SynCom, across all UV conditions, from independent replicate experiments. The x-axis represents the intensity of UVB radiation to which the plants were exposed. Low corresponds to 0.5 W $m^{-2}$, med to 1.0 W $m^{-2}$ and high to 2.2 W $m^{-2}$. Error bars represent standard error (n=8). **e,** *p*-values for the significance of the effect of listianol on the response of each strain to UV treatment across all three independent replicates.

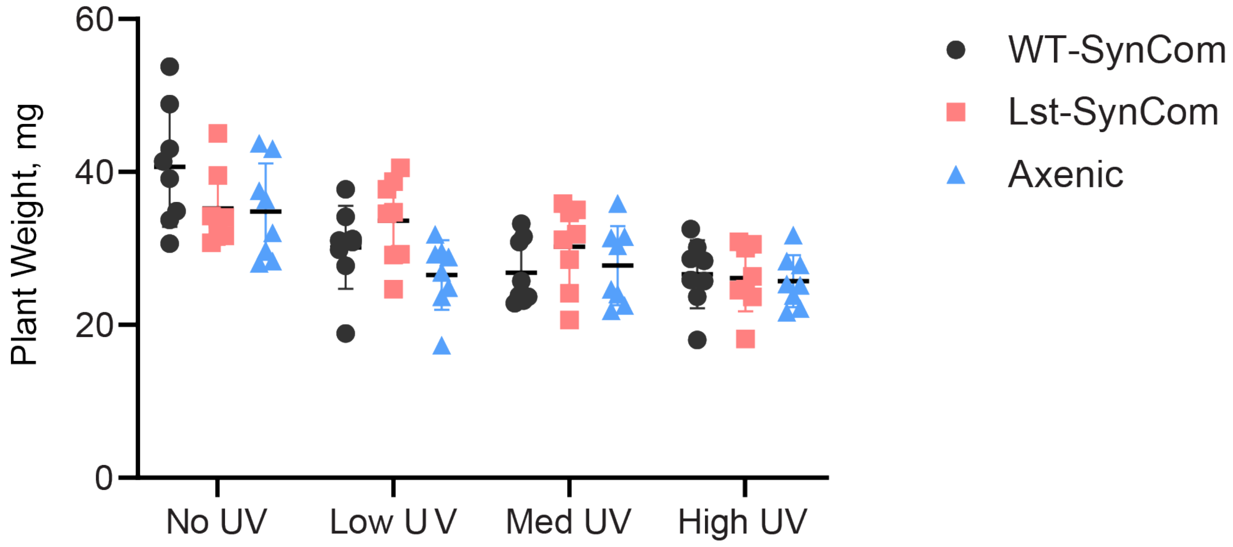

**Figure S17. Plant weight decreases with increasing intensity of UV treatment, but is not impacted by the presence or type of SynCom.** Weights (in mg) of plants inoculated with the indicated SynCom under increasing UV radiation intensity. Mean plant weight is indicated by the horizontal black bar and the vertical bars represent the standard deviation (n=8). A two-way ANOVA shows that plant weight is dependent on the presence of UV radiation ( p <0.0001), but does not depend on the presence or type of SynCom the plant was inoculated with (p = 0.0975).

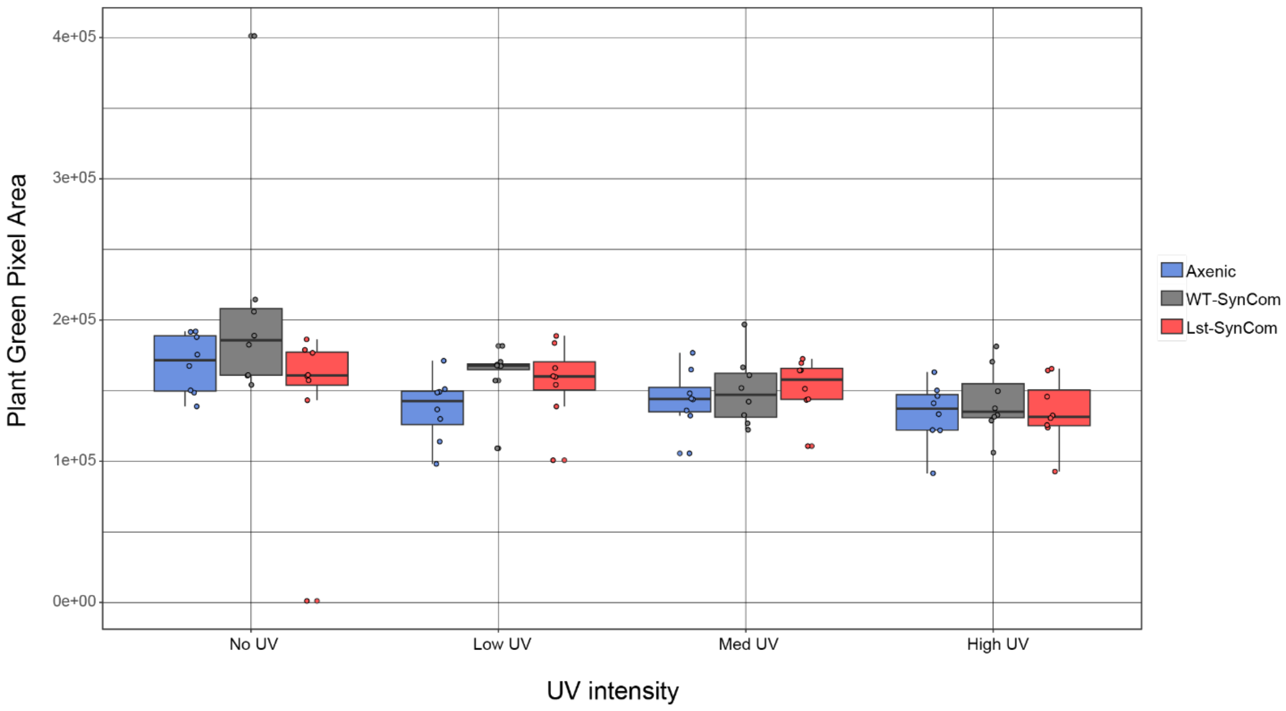

**Figure S18. Plant green pixel area deceases with UV treatment, but is not impacted by the presence or type of SynCom.** Plant green pixel area of plants inoculated with the indicated SynCom under increasing UV radiation intensity. The horizontal black line indicates the median, and the upper and lower edges of the box represent the 75th and 25th percentiles, respectively, while the whiskers indicate 1.5x interquartile range. A two-way ANOVA shows that green pixel area is dependent on the presence of UV ($p = 0.00621$), but not on the presence or type of SynCom the plant was inoculated with ($p = 0.06375$).

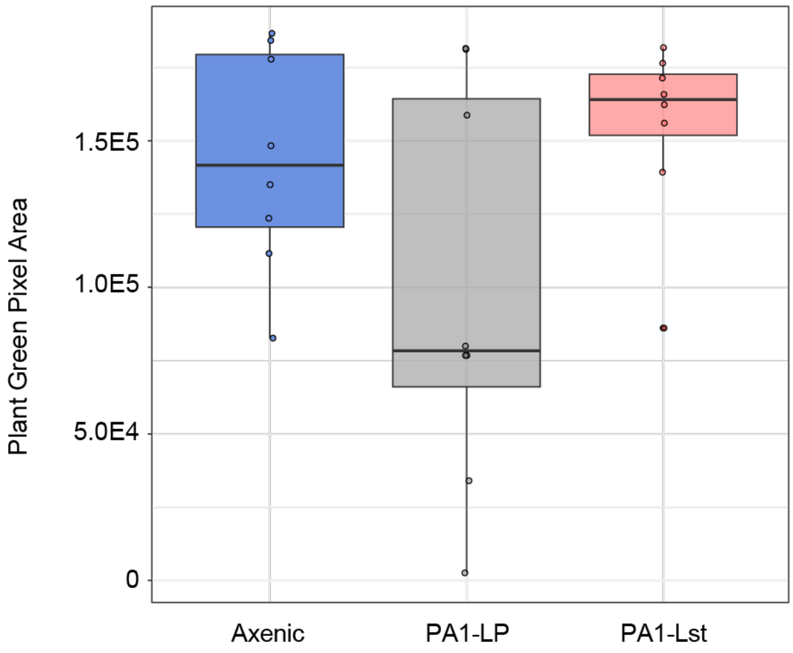

**Figure S19. Inoculation with PA1-Lst does not affect plant pigmentation.** Plant green pixel data 7 days post-inoculation with the indicated strains. The horizontal black line indicates the median, and the upper and lower edges of the box represent the 75th and 25th percentiles, respectively, while the whiskers indicate 1.5x interquartile range. Two-tailed unpaired t-tests (n=8) indicate no significant differences between axenic and PA1-Lst inoculated plants ($p = 0.526$) and PA1-LP and PA1-Lst inoculated plants ($p = 0.0513$).

### SUPPORTING TABLES

**Table S2. $^1$H (500 MHz) and $^{13}$C (125 MHz) NMR data assignment of listianol in DMSO-$d_6$ (J in Hz and $\delta$ in ppm) and $^1$H-$^1$H COSY and $^1$H-$^{13}$C HMBC correlations.**

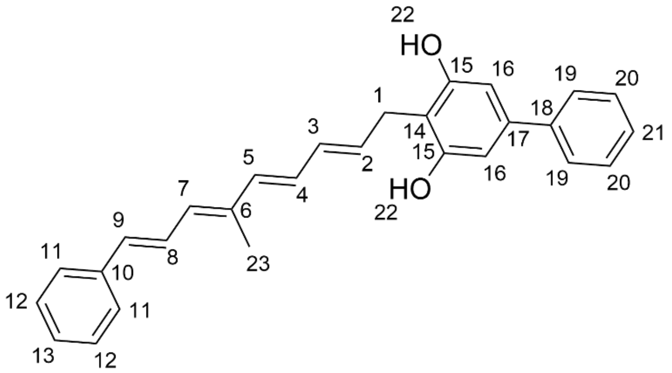

| Position | $\delta_C$, type | $\delta_H$, (type, J in Hz) | COSY | HMBC |
|---|---|---|---|---|
| 1 | 26.0, $CH_2$ | 3.34 (d, 6.8) | 2 | 2, 3, 14, 15 |
| 2 | 133.4, CH | 5.88 (dt, 15.0, 6.8) | 1, 3 | 1, 4, 14 |
| 3 | 130.2, CH | 6.12 (dd, 15.0, 10.5) | 2, 4 | 1, 4, 5 |
| 4 | 129.2, CH | 6.32 (dd, 15.0, 10.5) | 3, 5 | 2, 3, 5, 6 |
| 5 | 134.6, CH | 6.25 (d, 15.0) | 4 | 3, 4, 6, 7, 23 |
| 6 | 136.2, C | - | - | - |
| 7 | 130.8, CH | 6.25 (d, 11.3) | 8, 23 | 5, 8, 9, 23 |
| 8 | 125.6, CH | 7.24 (dd, 15.4, 11.3) | 7, 9, 12 | 6, 7, 9, 10 |
| 9 | 132.0, CH | 6.59 (d, 15.4) | 8, 12, 23 | 7, 8, 10, 11 |
| 10 | 137.3, C | - | - | - |
| 11 | 126.3, CH | 7.52 (d, m) | 12, 13 | 9, 13 |
| 12 | 128.6, CH | 7.32 (overlap) | 11, 13 | 10 |
| 13 | 127.3, CH | 7.21 (m) | 11, 12 | 11 |
| 14 | 111.9, C | - | - | - |
| 15 | 156.3, C | - | - | - |
| 16 | 104.7, CH | 6.56 (s) | - | 14, 15, 18 |
| 17 | 138.8, C | - | - | - |
| 18 | 140.6, C | - | - | - |
| 19 | 126.1, CH | 7.47 (d, m) | 20, 21 | 17, 21 |
| 20 | 128.8, CH | 7.42 (t, m) | 19, 21 | 18, 19, 21 |
| 21 | 127.0, CH | 7.32 (overlap) | 19, 20 | 18, 19 |
| 22 | - | 9.36 (s) | - | 14, 16 |
| 23 | 12.6, $CH_3$ | 1.94 (s) | 5, 7 | 5, 6, 7 |

**Table S3. Quantified inhibition of growth and carotenoid biosynthesis of diverse bacterial strains by listianol.**

| Strain | Growth inhibition MIC (µg/mL) | Carotenoid inhibition $IC_{50}$ (nM) |
|---|---|---|
| *Rathayibacter* sp. Leaf299 | 10 | >5,000 |
| *Rhizobium* sp. Leaf311 | >80 | NA |
| *Cellulomonas* sp. Leaf334 | 5 | 206 |
| *Chryseobacterium* sp. Leaf405 | 40 | NA |
| *Methylobacterium* sp. Leaf456 | >80 | 6.9 |
| *Methylorubrum extorquens* PA1 | >80 | 4.4 |
| *Methylobacterium* sp. 4-46 | >80 | 1.7 |
| *Clavibacter michiganensis* ssp. *michiganensis* 0317 | 5 | >5,000 |
| *Curtobacterium flaccumfaciens* B-729 | 10 | 352 |
| *Staphylococcus aureus* MN8 | 10 | 70.4 |
| *E. coli* BW25113 Δ*tolC* Δ*bamB* | >80 | NA |
| *Bacillus subtilis* PY79 | 40 | NA |
| *Erwinia carotovora* | 20 | NA |
| *Salmonella typhimurium* | >80 | NA |
| *Pseudomonas putida* | >80 | NA |
| *Enterococcus raffinosis* | >80 | NA |
| *Enterobacter aerogenes* | >80 | NA |
